# Astrocyte glutamate transport is modulated by motor learning and regulates neuronal correlations and movement encoding by motor cortex neurons

**DOI:** 10.1101/2022.01.20.477039

**Authors:** Chloe Delepine, Keji Li, Jennifer Shih, Pierre Gaudeaux, Mriganka Sur

## Abstract

While motor cortex is crucial for learning precise and reliable movements, whether and how astrocytes contribute to its plasticity and function during motor learning is unknown. Here we report that primary motor cortex (M1) astrocytes in mice show *in vivo* plasticity during learning of a lever push task, as revealed by transcriptomic and functional modifications. In particular, we observe changes in expression of glutamate transporter genes and increased coincidence of intracellular calcium events. Astrocyte-specific manipulations of M1 are sufficient to alter motor learning and execution, and neuronal population coding, in the same task. Mice expressing decreased levels of the astrocyte glutamate transporter GLT1 show impaired and variable movement trajectories. Mice with increased astrocyte Gq signaling show decreased performance rates, delayed response times and impaired trajectories, along with abnormally high levels of GLT1. In both groups of mice, M1 neurons have altered inter-neuronal correlations and impaired population representations of task parameters, including response time and movement trajectories. Thus, astrocytes have a specific role in coordinating M1 neuronal activity during motor learning, and control learned movement execution and dexterity through mechanisms that importantly include fine regulation of glutamate transport.

## INTRODUCTION

Astrocytes are now known to have diverse properties (Chai et al., 2017; Durkee and Araque, 2019; Khakh and Deneen, 2019; Khakh and Sofroniew, 2015; Martín et al., 2015; Slezak et al., 2019), and contribute in multiple ways to the modulation of brain information processing (Adamsky et al., 2018; Araque et al., 1999; Corkrum et al., 2020; Haydon, 2001; Hennes et al., 2020; Kol et al., 2020; Lines et al., 2020; Mederos et al., 2019; Nagai et al., 2019; Oliveira et al., 2015; Padmashri et al., 2015; Paukert et al., 2014; Perea et al., 2014a; Poskanzer and Molofsky, 2018; Poskanzer and Yuste, 2016; Santello et al., 2019; Sasaki et al., 2014; Yu et al., 2018). Previous studies have examined the role of astrocytes in learning and neuronal plasticity (Ackerman et al., 2021; Henneberger et al., 2010; Padmashri et al., 2015; Ribot et al., 2021; Suzuki et al., 2011). However, astrocyte contributions to neuronal population activity during behavioral tasks and learning remain largely unknown. While most studies of astrocyte-neuron interactions have been performed *in situ* in brain slices, a handful of *in vivo* studies have directly probed the effects of astrocytes on individual neurons (Perea et al., 2014b), or on neuronal populations (Lines et al., 2020; Poskanzer and Yuste, 2016; Yu et al., 2018). Here we investigated *in vivo* the role of cortical astrocytes in a motor learning task that specifically involves the coordinated activity of neuronal populations, where mice were rewarded for pushing a lever following an auditory cue (Peters et al., 2014). Primary motor cortex M1 has been implicated in motor learning (Peters et al., 2017; Tennant et al., 2011), including task acquisition (Gloor et al., 2015; Kawai et al., 2015; Nudo et al., 1996), and performance (Dombeck et al., 2009; Harrison et al., 2012; Peters et al., 2014), and we hypothesized that M1 astrocytes would have a role in mediating population neuronal activity and the task components mediated by the activity.

We found changes in M1 astrocyte gene expression during and after learning of the lever push task, with significant enrichment in glutamate transporters, suggesting a key role for astrocyte glutamate transport in primary motor cortex function during motor learning. We also found changes in M1 astrocyte calcium activity, with an increased coincidence of calcium events during the lever push associated with learning. Astrocytes influence synaptic transmission via glutamate transporters, and respond to, as well as modulate, neuronal activity through Gq-GPCR signaling (Adamsky et al., 2018; Agulhon et al., 2013; Aida et al., 2015; Lines et al., 2020; Rothstein et al., 1996). We thus reasoned that altering astrocyte glutamate transporters and Gq signaling would reveal effects of astrocytes on neuronal encoding and learned motor behavior. Glutamate transporter type 1 (GLT1) is prominently expressed in the cortex and hippocampus (Danbolt, 2001; Rothstein et al., 1994; Tanaka et al., 1997) and is almost exclusively found on astrocyte membranes, at the vicinity of synapses. Astrocytes express an extensive variety of G-protein coupled receptors (GPCRs) and second messenger systems to interact with and respond to the signals present in the extracellular environment (Kofuji and Araque, 2020; Porter and McCarthy, 1997). In particular, Gq-GPCR pathway activation in astrocytes of selected brain regions has been shown to influence specific behaviors and may be associated with modulation of glutamate transport (Adamsky et al., 2018; Agulhon et al., 2013; Cao et al., 2013; Chen et al., 2016; Lines et al., 2020; Martin-Fernandez et al., 2017; Scofield et al., 2015).

Using a transgenic mouse line in which we inhibited the expression of the glutamate transporter GLT1 locally in M1, we found that decreasing astrocyte glutamate clearance prevented learning of a stereotypical motor trajectory while preserving response time and task success rate. Expression of the engineered human muscarinic G protein-coupled receptor hM3Dq (Gq DREADD) in M1, activated by low doses of clozapine-N-oxide (CNO) (Agulhon et al., 2013; Armbruster et al., 2007; Roth, 2016), revealed that modulation of astrocyte Gq signaling impaired multiple parameters of task performance, leading to decreased performance rate, delayed response time and impaired learning of the trajectory. Using genetically encoded calcium indicators and high-resolution two-photon imaging, we imaged M1 layer 2/3 neuronal activity during execution of the motor task. Knockdown of astrocyte GLT1 increased the pool of active neurons and decreased neuronal correlations during movement. In contrast, activation of astrocyte Gq signaling abnormally increased neuronal correlations. Decoding and encoding models revealed changes in neuronal coding of task parameters following GLT1 reduction or Gq signaling increase, most critically in the representation of movement trajectories consistent with behavioral changes. These findings demonstrate specific *in vivo* contributions by astrocytes to the function of M1 layer 2/3 neuronal ensembles during motor learning, and their encoding of learned trajectories and task parameters.

## RESULTS

### Motor Learning Leads to Modification of Gene Expression Profiles in M1 Astrocytes

We trained mice to perform a cued lever push task (Peters et al., 2014) in which a lever press beyond a set threshold following trial start was rewarded with a water drop (Figure 1A). As expected, mice improved their success rate with training time, starting with a phase of rapid learning by day 3 (“novice” mice) and reaching a performance plateau after two weeks of training (“expert” mice) (Figure 1B).

**Figure 1:**
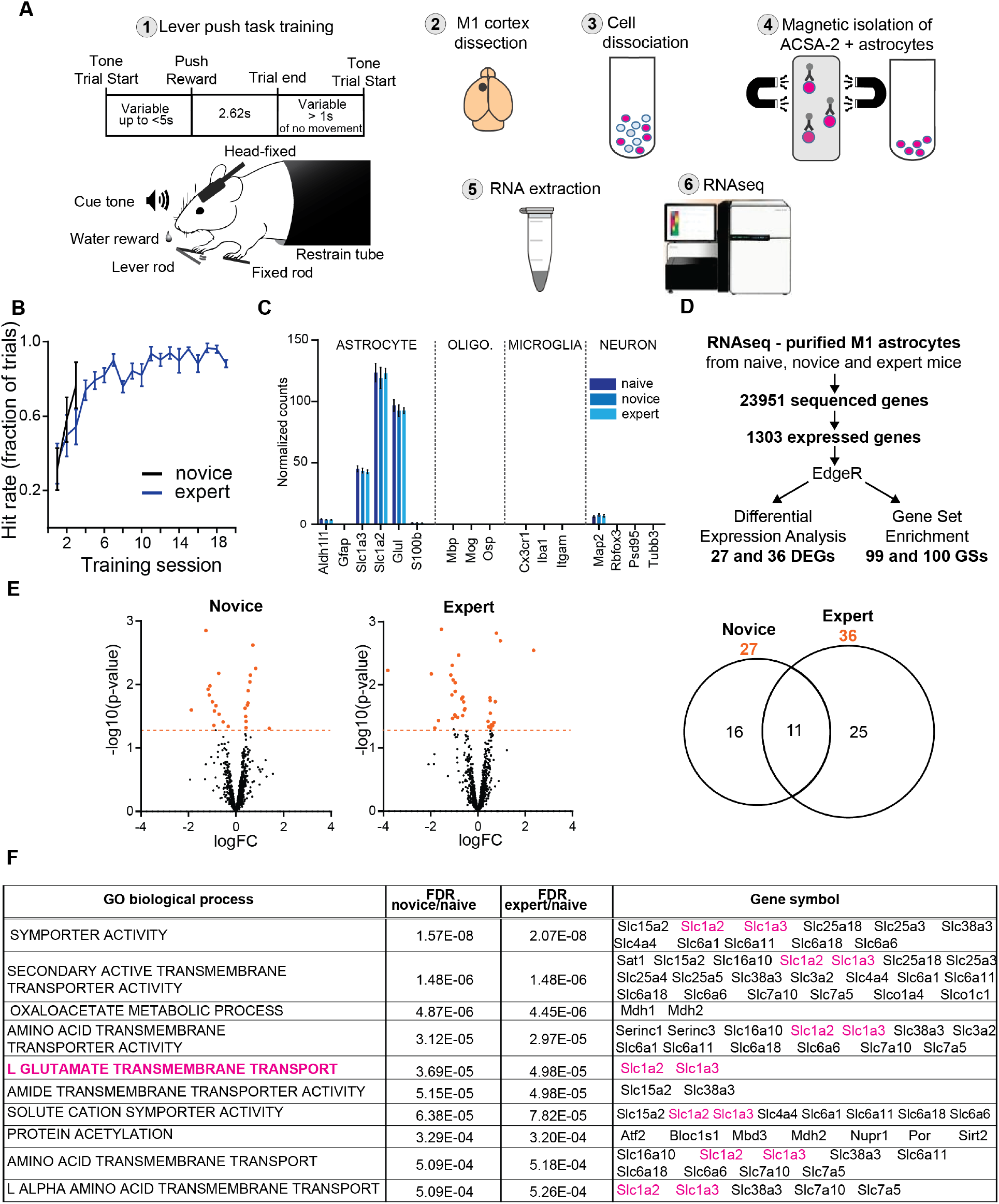
Motor Learning Leads to Modification of Gene Expression Profiles and Increased Coactivation of Microdomain Calcium Events in M1 Astrocytes. **A.** Methods summary. Lever push task: a tone indicated trial start; lever push within 5 secs was rewarded with a drop of water and response time and movement trajectory recorded. A window of 2.62 secs followed a push (correct trial) to allow consummatory licking of the reward, or signaled timeout (missed trial), after which a 1 sec inter-trial interval of no movement triggered start of the next trial. Motor cortex was dissected and cell dissociated. Astrocytes were trapped in a magnetic column using magnetic beads coated with anti-ACSA-2 antibodies and then collected for RNA extraction, cDNA library preparation and sequencing. **B.** Learning curves of novice and expert mice trained in the lever push task. Groups were named as follows: naive mice, not trained in the lever push task; novice mice, trained for 3 training sessions; and expert mice, trained for 19 training sessions and showing successful learning of the task. Graph represents hit rate (mean ± SEM) as measured by the fraction of correct trials. n=6 wildtype mice for each group. **C.** Astrocyte purification was confirmed for the three groups by measures of the normalized gene expression of astrocyte, oligodendrocyte (oligo.), microglia and neuron specific genes. Gene expression was normalized by housekeeping gene counts (*Gapdh*). Bar plots represent mean ±SEM. **D.** Gene expression profiles from astrocytes of the three groups were analyzed and compared using the Bioconductor’s EdgeR package to perform Differentially Expressed Genes (DEGs) and Gene Set Enrichment Analyses. 27 DEGs were identified in novice mice, 36 DEGs in expert mice (see Supplemental Figure 1 and **Supplemental Table 1**). 99 gene sets in novice mice and 100 in expert mice were significantly enriched relative to naive mice, with an overlap of 98 Gene Ontology (GO) categories (see **Supplemental Table 3**). N=6 wildtype mice for each of the 3 groups. **E.** We identified 27 Differential Expression Genes (DEGs) in naïve mice, 36 DEGs in expert mice; 11 DEGs were common for both. **Left:** Volcano plots of logarithms of fold change (log2FC) and p-value (-log10 (p-value)) of differential expression of all expressed genes. Each dot represents one gene. Orange dots indicate DEGs (p-value>0.05) **Right:** Venn diagram of DEGs. **F.** Top 10 significantly enriched Gene Sets differentially regulated in M1 astrocytes in novice and expert mice compared to naive mice, and their respective expressed genes. In particular, the gene set corresponding to L-glutamate transport function, in blue, was significantly enriched.

We first used RNA sequencing (RNAseq) to identify gene expression changes in M1 astrocytes associated with learning. M1 cortices of mice were extracted after no training (untrained “naïve” mice), training in the lever push task for three days (partially trained novice mice), or training in the lever push task for nineteen days (fully trained expert mice) (Figure 1A). To match stress levels, all three groups were water-restricted and head-fixed for the same duration as the expert mice. Astrocytes were isolated using ACSA-2 immuno-magnetic sorting (Holt and Olsen, 2016) (Figure 1A). We validated the isolation protocol by comparing the normalized gene counts of cell-type specific markers for the three groups. Samples of all the groups were similarly enriched in astrocyte-specific genes and depleted of other brain cell markers (Figure 1B). RNAseq was performed and results analyzed using the EdgeR package (Bioconductor) to identify (1) differentially expressed genes (DEGs) and (2) significantly enriched gene sets in astrocytes from novice and expert mice compared to naive mice (Figure 1D). We found 27 DEGs in novice mice and 36 DEGs in expert mice (p-value < 0.05) with an overlap of 11 DEGs (Figure 1E, **Supplemental Table 1**). The numbers of DEGs which were up or down-regulated were similar (Figure 1E). We used the PANTHER classification system to analyze the DEG list (Mi et al., 2019). Several Gene Ontology (GO) biological processes and molecular functions were enriched in the DEG list of known protein coding genes (**Supplemental Table 2**). The differentially regulated GO categories were mostly related to metabolism, transcription and signaling. Moreover, DEGs were significantly enriched in membrane or extracellular protein coding genes, suggesting the importance of transporters, receptors and cell-cell communication (**Supplemental Table 2**).

We performed a Gene Set Enrichment Analysis (GSEA) of the RNAseq data, which identifies sets of genes with expression changes that may be small, and therefore not identified individually by DEG analysis, but that collectively contribute to the dysregulation of a shared biological function (GO category). We identified 99 and 100 gene sets in novice and expert mice respectively, that were significantly enriched relative to other genes in terms of differential expression (Figure 1D). Most of the enriched gene sets were overlapping GO categories of transmembrane transporters such as Symporter Activity, Secondary Active Transmembrane Transporter Activity, Amino Acid Transporter Activity, L-Glutamate Transmembrane Transport, Organic Acid Transmembrane Transport, and Sodium Ion Transmembrane Transporter (Figure 1F, **Supplemental Table 3**). The GO: L-Glutamate Transport gene set contained only two genes, coding for the two astrocyte-specific glutamate transporters GLT1 (*Slc1a2*) and GLAST (*Slc1a3*), with high and low cortical expression levels respectively. Not only was this gene set significantly enriched, it was also included in most of the other enriched GO category sets (Figure 1F, **Supplemental Table 3**). The GO: Solute Sodium Symporter Activity gene set was also contained in most of the enriched gene sets, with 6 genes expressed in M1 cortex samples including astrocyte-specific glutamate transporters (*Slc1a2*/GLT1 and *Slc1a3*/GLAST) and GABA transporters (*Slc6a1*/GAT1 and *Slc6a11*/GAT3) (**Supplemental Table 3**). We validated with RTqPCR the expression changes of genes in these gene sets in expert WT mice compared to naive WT mice (Supplemental Figure 1A). To control for the specificity of the observed changes to the forelimb M1 cortex, we performed the same experiment with left hindlimb M1 (hM1) cortex samples (Supplemental Figure 1B). We observed no significant differences in hM1 expression levels for most genes, but we noted a trend for a few genes, and a significant change for one gene, *Slc6a1*, to be upregulated in expert mice in both the forelimb and hindlimb motor cortex, supporting the idea that large regions of mouse M1 and even wider swaths of cortex are partially activated during reward-related movement (Musall et al., 2019).

These results indicate that M1 cortical astrocytes undergo changes in gene expression associated with motor learning that may underlie mechanisms of astrocyte contributions to M1 function. Furthermore, they highlight the importance of glutamate transporter modulation for our motor learning task.

### Motor Learning Leads to Increased Coactivation of Calcium Events in M1 Astrocyte

In addition to changes in gene expression, we assessed functional changes in astrocytes as a result of learning the lever push task. Astrocytes respond to neuronal and synaptic activity with complex spatiotemporal fluctuations in intracellular calcium, localized in their soma, branches and in “microdomains” within their fine processes (Agarwal et al., 2017; Arizono et al., 2020; Bindocci et al., 2017; Di Castro et al., 2011; Shigetomi et al., 2013; Srinivasan et al., 2015; Stobart et al., 2018). To determine if astrocyte calcium signals reflect motor learning, we performed chronic imaging of calcium activity using the membrane-bound calcium indicator GCaMP6f-Lck driven by the Aldh1l1 promoter, and detected and analyzed calcium events using the AQuA algorithm (Wang et al., 2019) (Figure 2A, **Supplemental Video 1**). After two weeks of training, mice showed increased coincidence of calcium activity during the lever push (Figure 2). In novice animals, the calcium events detected during the lever push were rarely coincident, moreover their DF/F0 had low cross-correlations, indicating that the fluorescence changes detected as movement-related events were not z-movement artifacts (Supplemental Figure 2). While the number, area and amplitude of individual events did not change in expert vs. novice mice (novice=0.9741±0.074 DF/F0, expert=0.9559±0.053 DF/F0, n=7) (Figure 2B), the percentage of trials in which two or more events occurred concurrently during the movement was increased (novice=0.15±0.052, expert=0.2146±0.043, n=7) (Figure 2C-D). As a consequence, the average trial activity was increased (novice=0.1375±0.01, expert=0.1837±0.02, n=7) (Figure 2E-F). These results indicate that the pattern of calcium signaling in M1 astrocytes changes during the course of motor learning, potentially via the regulation of localized calcium events by GLT1 (see Discussion). This enhanced coincidence of astrocyte calcium events could reflect, as well as contribute to, the coordinated activation of M1 synapses and neurons.

**Figure 2:**
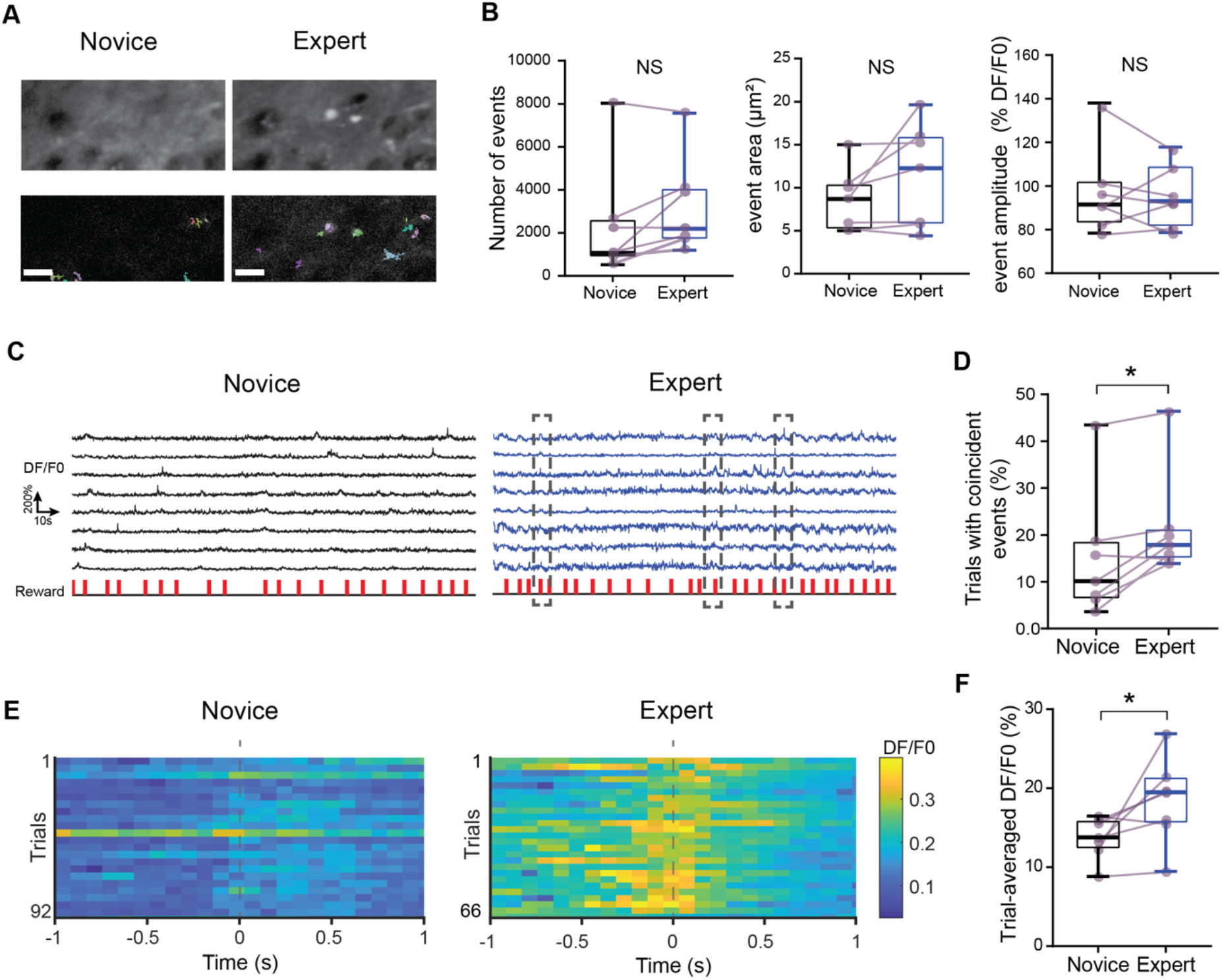
Motor Learning Leads to Increased Coactivation of Calcium Events in M1 Astrocyte. GCaMP6f-lck signals in astrocyte were imaged in layer 2/3 of M1 as wild-type mice performed the lever push task. For each mouse, spatiotemporal calcium events were detected from 3 non-overlapping field of views, during 3 novice and 3 expert sessions. N= 7 mice. **A-B.** Characterization of detected astrocyte calcium spatiotemporal events. **A.** Example frames from early in training (Novice) compared with an expert session of training (Expert) are shown, with astrocytic calcium events identified by the AQuA algorithm shown in the overlay (bottom panels). Scale bar represents 10μm**. B.** Quantification of number, area, and amplitude (DF/F0) of the detected astrocyte calcium events from novice and expert training sessions. Number of detected events: mean = 2331±992.7 (median=1062) events for novice, 3199±837.4 (median=2190) for expert mice, NS: p=0.1094, paired Wilcoxon test, n=7. Event area: mean = 8.602±1.368 µm^2^ (median= 8.701 µm^2^) for novice, 11.32±2.258 µm^2^ (median=12.27 µm^2^) for expert mice, NS: p=0.0781, paired Wilcoxon test, n=7. Amplitude: mean=0.9741±0.074 DF/F0 (median=0.9151 DF/F0) for novice, 0.9559±0.053 DF/F0 (median=0.9304 DF/F0) for expert mice, NS: p=0.6875, paired Wilcoxon test, n=7. Box plot bar represents median, box extends from the 25th to 75th percentile, and whiskers show 10th to the 90th percentile. Purple circles represent the data points for each individual mouse. Paired values (same mouse) are indicated with purple lines. **C-F**. Learning of the task was associated with an increase in coincident activity of astrocyte calcium events **C.** DF/F0 traces of example astrocytic calcium events. Red bars below DF/F0 traces indicate reward time. Boxes in the expert traces indicate coincident events. **D.** Percentage of trials where there was coincident activity (2 or more events occurring at the same time) during the lever push, in novice training sessions compared to expert training sessions. Novice: mean=0.15±0.052 (median=10.12%), expert: mean=0.2146±0.043 (median=17.87%) *: p=0.0313, paired Wilcoxon test, n=7. **E.** Trial-averaged astrocyte calcium activity (DF/F0) in example novice and expert sessions from the same M1 layer 2/3 astrocytes. Calcium activity in astrocytes was measured during the task and events were identified, aligned to the threshold crossing in the movement trajectory, and averaged. Zero (0) on the time axis and the vertical dashed line indicate time when lever position reached the reward threshold (1mm). **F.** Quantification of astrocyte calcium activity during correct trials (trial-averaged DF/F0) in novice and expert mice. Novice: mean=0.1375±0.01 (median=0.1378 DF/F0), expert: mean=0.1837±0.02 (median=0.195 DF/F0). *: p=0.469, paired Wilcoxon test.

### Decreased GLT1 Levels and Astrocyte Gq Pathway Activation in M1 Impair Motor Learning-Associated Changes in Gene Expression

Our gene expression analyses strongly implicated the astrocyte glutamate transporter GLT1 in learning-related changes. GLT1 critically influences synaptic transmission, as shown *in vitro* and in slices *in situ* (Aida et al., 2015; Arnth-Jensen et al., 2002; Cui et al., 2014; Huang et al., 2004; Oliet et al., 2001; Omrani et al., 2009; Rothstein et al., 1996; Takayasu et al., 2006; Tanaka et al., 1997; Tsukada et al., 2005; Tzingounis and Wadiche, 2007). Moreover, astrocyte GLT1 has an important role in neuronal plasticity, as demonstrated *in situ* (Filosa et al., 2009; Oliet et al., 2001; Omrani et al., 2009). While mouse models of GLT1 knockdown have shown major behavioral deficits, previous studies largely involved brain-wide and complete knockdown (Aida et al., 2015; Cui et al., 2014; Gomez et al., 2019; Niederberger et al., 2003; Pardo et al., 2006). To specifically explore the role of GLT1 expression in M1 astrocytes *in vivo* during motor learning, we delivered a viral vector encoding the CRE-recombinase under the astrocyte-specific GFAP promoter unilaterally in M1 cortex of GLT1-flox heterozygous mice (“GLT1”) and their wild-type littermates (“WT”) (Figure 3A). Two weeks after injection, the expression level of GLT1 was decreased to around 50% at both mRNA and protein levels (mRNA ratio=0.54 ± 0.0.088, WT n=10, GLT1 n=6; protein ratio=0.55 ± 0.049, WT n=6, GLT1 n=8) (Figure 3B, C; Supplemental Figure 3).

**Figure 3:**
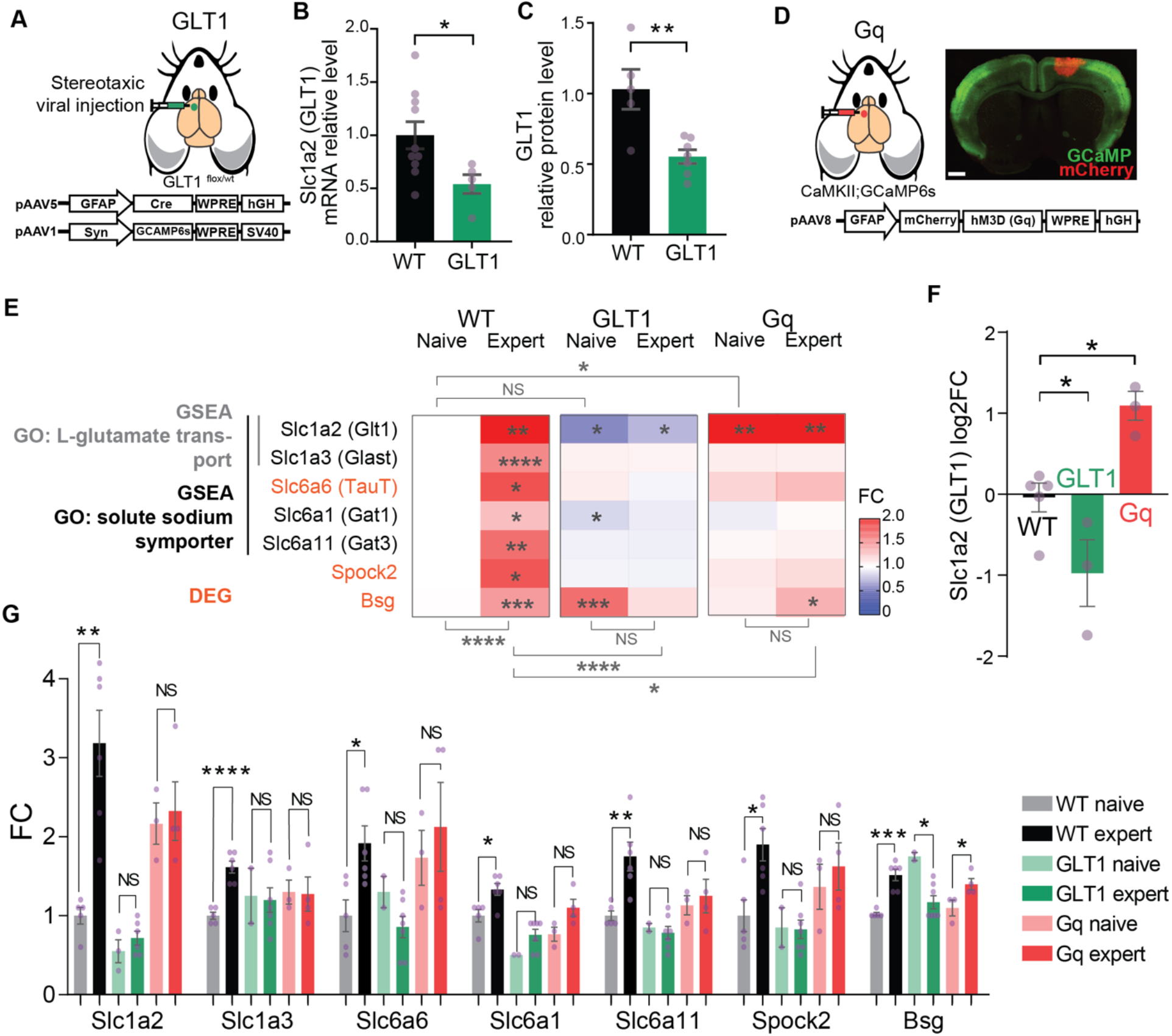
Decreased GLT1 Levels and Astrocyte Gq Pathway Activation in M1 Impair Motor Learning-Associated Changes in Gene Expression. **A.** AAV-GFAP-CRE, or alternatively AAV-GFAP-CRE and AAV-Syn-GCAMP6s (for neuronal imaging experiments), were injected in M1 of GLT1 flox/+ mice (“GLT1”) and their wild type littermates (“WT”). **B.** GLT1 mice show a 46% reduction of Slc1a2 (Glt1) mRNA levels compared to WT (n= 10 WT mice, 6 GLT1 mice, ratio=0.5401±0.0882, *: p=0.0337, unpaired t-test), as measured by RTqPCR. Bar plots in B, C represent mean ± SEM, purple dots represent single observations. **C**. GLT1 mice show a 45% reduction of GLT1 protein level (n=6 WT, 8 GLT1 mice, ratio=0.5535±0.04859, **: p=0.0045, unpaired t-test), as measured by Western Blot. See also Supplemental Figure 2**. D. Left:** AAV-GFAP-h3MD(Gq)-mCherry was injected in the M1 cortex of CaMKII-GCaMP6s or WT mice. **Right:** Example CaMKII-GCAMP6s mouse brain coronal section. AAV-GFAP-h3MD(Gq)-mCherry expression was localized in upper layer astrocytes of M1. Scale bar, 1mm. **E.** Heatmap of average gene expression fold change (FC) of selected genes, measured by RTqPCR and normalized to wildtype naïve mice. Genes were selected within RNAseq identified DEGs (*Bsg*, *Spock2* and *Slc6a6*, highlighted in orange) and GSEA gene sets (*Slc1a2*, *Slc1a3* from GO: L-glutamate transmembrane transport, and *Slc1a2*, *Slc1a3*, *Slc6a6*, *Slc6a1*, *Slc6a11* from GO: solute sodium symporter activity) (see also Figure 1). In contrast to WT mice, GLT1 and Gq mice did not show motor learning-associated changes in gene expression. Naïve vs Expert, two-way ANOVA, WT: ****: p<0.0001, GLT1: NS: p=0.2338, Gq: NS: p=0.1603 for the variability explained by learning. Stars on the heatmap indicate statistically significant differences for each individual gene in comparison to WT naïve (one-way ANOVA with Dunnett’s multiple comparisons test), stars below and above the heatmap indicate statistical significance of the variability explained by the manipulation considering all genes (two-way ANOVA), stars over the bar plot indicate statistical significance for the mean comparison of naïve and expert expression levels for each individual gene. NS: p>0.05, *: p<0.05, **: p<0.01, ***: p<0.001, ****: p<0.0001. **F.** *Slc1a2* was significantly downregulated in GLT1 naïve mice and upregulated in Gq naïve mice compared to WT naïve mice. Logarithm of fold change (log2FC). Naïve WT mice: mean=0±0.1792, n=5; naïve GLT1 mice, mean=-0.9738±0.4123, n=3, Gq naïve: mean=1.09±0.1783, n=3. WT vs. GLT1 *: p=0.0197, WT vs. Gq *: p= 0.071, Dunnett’s multiple comparisons test, One-way ANOVA). **G.** Bar plot (mean± SEM) representing the expression fold change (FC) of selected genes in the forelimb motor cortex of WT, GLT1 and Gq mice, naïve and expert mice. RTqPCR confirmed significantly increased expression levels of all selected genes in expert WT mice compared to naïve WT mice (n=5 naïve WT mice, n= 6 expert WT mice). Compared to WT naïve mice, *Slc1a2* was significantly downregulated in GLT1 naïve and expert mice (WT naïve, mean=1±0.1049, n=5; GLT1 naïve, mean=0.55±0.1443, n=3, *: p=0.0424, unpaired t-test; GLT1 expert, mean=0.7167±0.08724, n=5; *: p=0.0476, Mann Whitney U test), and upregulated in Gq naïve and expert mice (Gq naïve : mean=2.167±0.2603, n=3,**: p=0.0026, unpaired t-test; Gq expert : mean=2.325±0.3705, n=4, **: p=0.0465, unpaired t-test). *Bsg* was the only gene that showed changes between naïve and expert mice in GLT1 and Gq mice, with a downregulation associated with the learning in GLT1 mouse and an upregulation in Gq mice (GLT1: naïve, mean=1.75±0.05, n=3, expert, mean=1.171±0.08371, n= 7, *: p=0.0101, unpaired t-test; Gq: naïve, mean =1.1±0.1, n=3, expert, mean=1.40±0.07071, n=4, *: p=0.0286, Mann Whitney U test). Of note, *Bsg* was significantly upregulated in naïve GLT1 mice and in expert Gq mice compared to WT naïve mice (GLT1 naïve: mean = 1.75±0.05, n=3, ****: p<0.0001, unpaired t-test; Gq expert: mean = 1.40±0.07071, n=4, ***: p=0.0007, unpaired t-test).

Gq pathway activation in astrocytes has diverse effects on astrocytes, affecting calcium release from intracellular stores, astrocyte-neuron functions and specific behaviors (Adamsky et al., 2018; Agulhon et al., 2013; Cao et al., 2013; Chen et al., 2016; Martin-Fernandez et al., 2017; Scofield et al., 2015). To explore the effect of the Gq pathway in M1 astrocytes, we used an engineered Gq-coupled designer receptor (DREADD) hM3Dq that can be activated by exogenous clozapine-N-oxide (CNO) in a time-restricted manner (Armbruster et al., 2007; Roth, 2016). We injected a viral hM3Dq-mCherry construct unilaterally in M1 under the astrocyte specific GFAP promoter (Figure 3D) (Armbruster et al., 2007; Roth, 2016). Co-staining with astrocyte marker S100beta showed high specificity and high density of expressing cells (>98%) (Supplemental Figure 4A).

Based on the motor learning-associated changes in transcriptomic expression of genes and gene sets in M1 astrocytes of wildtype mice (Figure 1), we explored the expression of a selection of these genes using RTqPCR in Gq and GLT1 naïve and expert mice (Figure 3E-G). Genes were selected within the previously identified DEGs (*Bsg*, *Spock2* and *Slc6a6*) and GSEA gene sets (*Slc1a2*, *Slc1a3* from GO: L-glutamate transmembrane transport, and *Slc1a2*, *Slc1a3*, *Slc6a6*, *Slc6a1*, *Slc6a11* from GO: solute sodium symporter activity) (Figure 3E). First, all of the genes showed significant upregulation in expert WT mice compared to untrained (naïve) WT mice, confirming the RNAseq results in independent samples and experiments (Figure 3E). In contrast, both GLT1 and Gq mice showed no learning-associated difference for the selected genes, with the exception of *Bsg* which was significantly downregulated in GLT1 expert mice compared to GLT1 naïve mice and significantly upregulated in Gq expert mice compared to Gq naïve mice (Figure 3E, G). *Slc1a2* (GLT1) was significantly downregulated in GLT1 mice as expected (Figure 3B, C; F, G), but significantly upregulated in Gq mice (Figure 3F, G).

Thus, Gq activation and GLT1 reduction both impair motor learning-associated gene expression changes, and have opposite effects on GLT1 expression. These analyses led us to examine behavioral and neuronal consequences of GLT1 reduction and Gq activation during motor leaning.

### Decreased GLT1 Levels in M1 Astrocytes Alter Movement Trajectories

In mice trained daily in the lever push task (Figure 1A), WT mice improved their success rate and decreased their response time with training (Figure 4A, B), as previously shown (Peters et al., 2014). Additionally, their lever push movements became more stereotyped (more similar across trials) and more precise (smoother) (Figure 4C-E). GLT1 mice had similar success rates and response times as WT controls (Figure 2A, B). However, GLT1 mice showed deficits in learning-associated stereotyped movements, as indicated at training days 12-14 by the reduced average pairwise trial-to-trial similarity of the movement trajectory (average pairwise correlation WT 0.82±0.011 n=15, GLT1 0.71±0.014 n=12) and low dexterity (smoothness coefficient WT 0.73±0.031 n=15, GLT1 0.46±0.032 n=12) (Figure 2F-H). Thus, a reduction of astrocyte GLT1 expression in M1 is sufficient to perturb the stereotypy and smoothness of movement trajectories that accompany motor learning.

**Figure 4:**
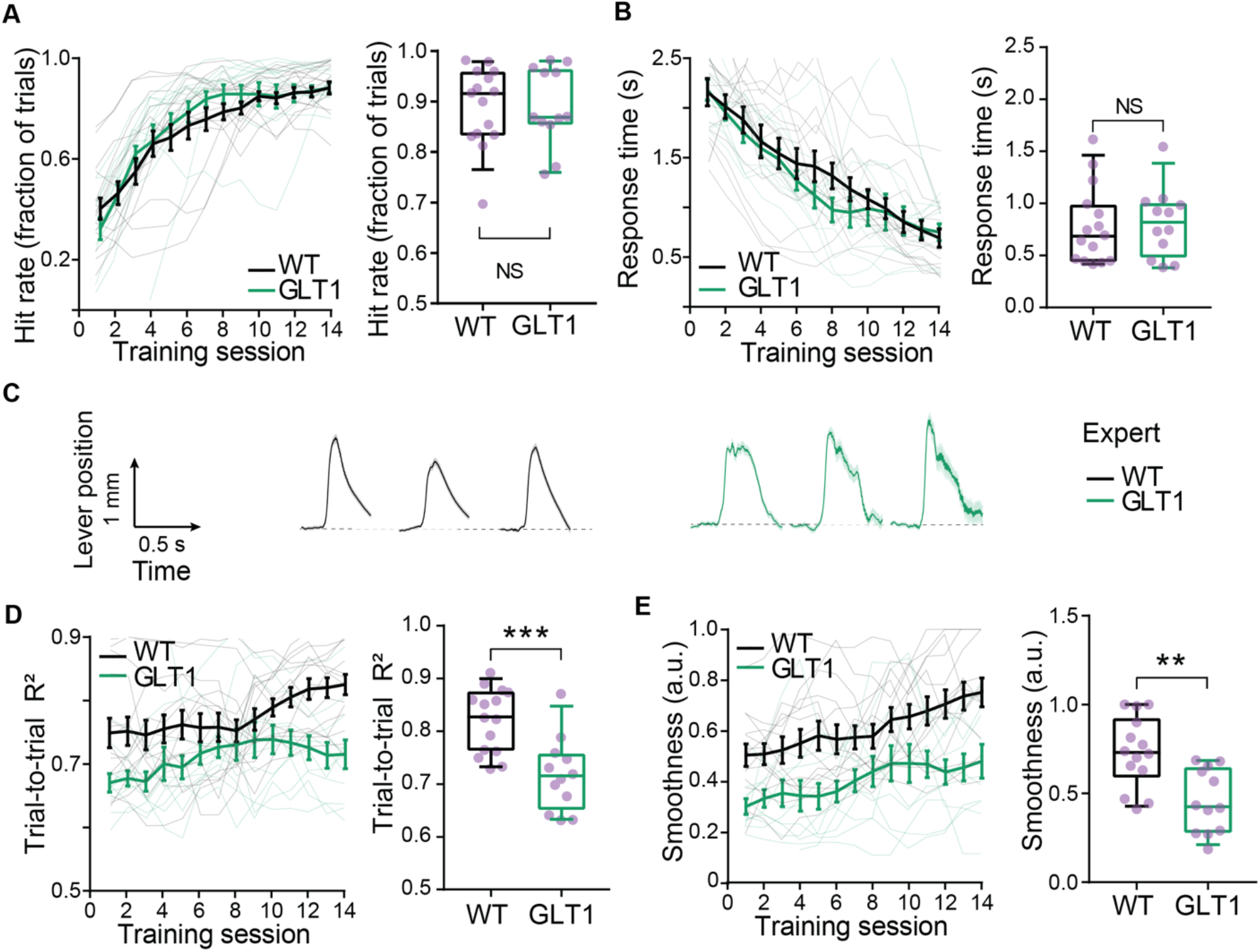
Decreased GLT1 Levels in M1 Astrocytes Alter Movement Trajectories. **A.** GLT1 reduction in M1 astrocytes had no effect on hit rate. **Left:** all training sessions **Right:** average of expert sessions (training days 12-14) (WT: mean=0.8892±0.01968, GLT1: mean=0.8847±0.01774, NS: p= 0.984, unpaired t-test). For all panels, N= 15 WT, 12 GLT1 mice; box plots as described in Fig. 2B. **B**. GLT1 reduction had no effect on response time. **Left:** all training sessions **Right:** average of expert sessions (WT: mean=0.7739±0.0682s, GLT1: mean=0.7905±0.06986, NS: p= 0.8384, unpaired t-test). **C-E.** Reduced GLT1 in M1 astrocytes perturbed learning of stereotyped and smooth movement trajectories **C.** Example average lever trajectory traces of three expert training sessions for one WT (black) and one GLT1 (green) example mouse. **D.** Trial-to-trial movement similarity estimated by the average pairwise correlation of the movement traces (trial-to-trial R^2^). **Left:** all training sessions. **Right:** expert sessions average (WT: mean WT=0.8204±0.01064, GLT1: mean=0.7137±0.0139; ***: p=0.0004, unpaired t-test)**. E.** Average movement smoothness estimated by the inverse of the number of push events per movement. **Left:** all training sessions. **Right:** expert sessions average (WT: mean=0.7325±0.03158, GLT1: mean=0.4585±0.03156; **: p=0.0012, unpaired t-test).

### Astrocyte Gq Pathway Activation in M1 Impairs Task Performance

Activation of Gq-DREADD expressing astrocytes upon CNO application has been shown in brain slices to induce a release of calcium from the intracellular stores and an increase of intracellular calcium signaling (Agulhon et al., 2013). However, to our knowledge, this effect has rarely been examined *in vivo.* We injected unilaterally in M1 a viral GFAP-hM3Dq-mCherry construct in astrocyte-specific cytoplasmic GCaMP expressing mice (GFAP-GCaMP5G mouse line) to image *in vivo* calcium activity in astrocytes, 30 min after intraperitoneal injection of CNO. We observed that a low dose of CNO triggered an increase in intracellular calcium as measured by the baseline GCaMP fluorescence (Supplemental Figure 4B), along with a decrease in frequency and amplitude of calcium events (Supplemental Figure 4C-E), consistent with near saturation of signaling due to the depletion of internal calcium stores. Similarly, a recent study found that Gq-DREADD activation in cortical astrocytes almost completely abolished calcium dynamics (Vaidyanathan et al., 2021).

We trained Gq-DREADD expressing mice (“Gq”) and controls (“CTRL”) in the lever push task. Two weeks after virus injection, mice were trained daily for 14 days of training sessions, with an IP injection of a low dose of CNO 30 min before training started (Figure 5). Training was continued for six additional days with an injection of vehicle (saline) solution instead of CNO (Figure 5). Gq mice injected with CNO showed a decreased performance rate (average hit rate Gq+CNO 0.64±0.038 n=7, CTRL+CNO 0.81±0.029 n=13), as measured by the fraction of successful trials, that improved rapidly after withdrawal of CNO (saline injection) (average hit rate Gq+saline 0.82±0.032 n=7) (Figure 5A). Gq mice injected with CNO also had increased response times (average response time, Gq+CNO 1.46±0.10 n=7, CTRL+CNO 0.71±0.077 n=13) that improved upon CNO withdrawal (average response time, Gq+saline 1.02±0.13 n=7) (Figure 5B). Finally, Gq mice showed a decreased stereotypy of movement, as indicated by the lower average pairwise trial-to-trial similarity of the movement trajectories (average pairwise correlation CTRL+CNO 0.73±0.013 n=13, Gq+CNO 0.68±0.016 n=7). This was rescued by withdrawal of CNO (average pairwise correlation, Gq+saline 0.75±0.016 n=7) (Figure 5D). However, we did not observe any significant difference in movement smoothness (Figure 5E). Thus, Gq signaling activation in M1 astrocytes during motor learning is sufficient to temporarily perturb task performance by decreasing performance rate, slowing responses and reducing the stereotypy of movement trajectories.

**Figure 5:**
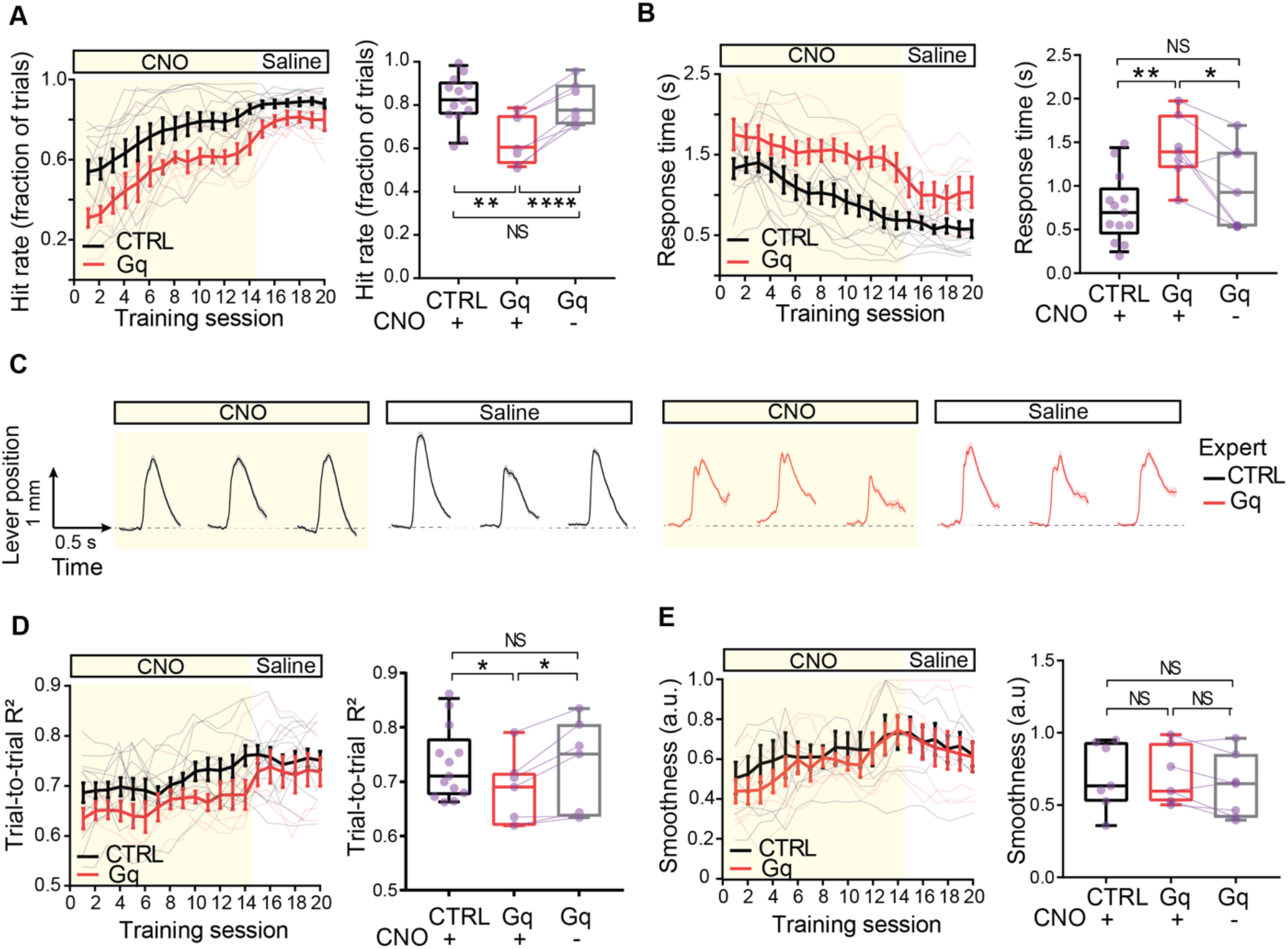
Astrocyte Gq Pathway Activation in M1 Impairs Task Performance. Mice expressing GFAP-h3MD(Gq)-mCherry (“Gq”) and controls (“CTRL”) were injected intraperitoneally 30 min before a training session started with low dose of clozapine-N-oxide (“CNO”) for the first 14 training days, then with saline solution (“saline”) for 6 additional training days. For all panels, N=13 CTRL, 7 Gq mice; box plots as described in Fig. 2B. **A.** Gq activation in M1 astrocytes reduced hit rate. **Left:** all training sessions **Right:** average of expert sessions (data from training days 12-14 and 18-20) (CTRL+CNO: mean=0.8127±0.02985, Gq+CNO: mean=0.6445±0.03813, Gq+saline: mean=0.8205±0.03272, CTRL+CNO vs. Gq+CNO **: p=0.0062, CTRL+CNO vs. Gq+saline NS: p=0.9660, Tukey’s multiple comparison, One-way ANOVA; Gq+CNO vs. Gq+saline ****: p<0.0001, paired t-test). **B**. Gq activation increased response time. **Left:** all training sessions **Right:** average of expert sessions (CTRL+CNO: mean=0.7104±0.07735, Gq+CNO: mean=1.458±0.1039, Gq+saline: mean=1.02±0.1249, CTRL+CNO vs. Gq+CNO **: p=0.0048, CTRL+CNO vs. Gq+saline NS: p=0.3961, Tukey’s multiple comparison, One-way ANOVA; Gq+CNO vs. Gq+saline *: p=0.0225, paired t-test). **C-E.** Gq activation in M1 astrocytes perturbed movement trajectories. **C.** Example average movement trace of three expert training sessions with CNO or saline injection, for one CTRL (black) and one Gq (red) example mouse. **D.** Trial-to-trial movement similarity. **Left:** all training sessions **Right:** average of expert sessions (CTRL+CNO: mean=0.73±0.01295, Gq+CNO: mean=0.681±0.01631, Gq+saline: mean=0.7467±0.01579, CTRL+CNO vs. Gq+CNO *: p=0.0493, CTRL+CNO vs. Gq+saline NS: p=0.9668, Tukey’s multiple comparison, One-way ANOVA; Gq+CNO vs. Gq+saline *: p=0.0111, paired t-test). **H** Average movement smoothness **Left:** all training sessions. **Right:** average of expert sessions (CTRL+CNO: mean=0.6999±0.04929, Gq+CNO: mean=0.7005±0.04288, Gq+saline: mean=0.6999±0.04929, CTRL+CNO vs. Gq+CNO NS: p>0.999, CTRL+CNO vs. Gq+saline NS: p=0.8073, Tukey’s multiple comparison, One-way ANOVA; Gq+CNO vs. Gq+saline NS: p=0.0855, paired t-test).

As hit rate and response time were not fully rescued by CNO withdrawal, we examined a CNO-independent effect of Gq-DREADD by injecting Gq mice with saline solution, 30 min before training, throughout the training. We did not observe any significant difference between the control groups (Supplemental Figure 5A, B). Thus, the residual effects on behavioral performance in Gq-DREADD mice treated with CNO are likely the result of disrupted Gq signaling in astrocytes.

### Decreased GLT1 Levels in M1 Astrocytes Reduce Neuronal Signal Correlations

GLT1 knockdown has been shown to drive neuronal hyperexcitability (Aida et al., 2015; Arnth-Jensen et al., 2002; Filosa et al., 2009; Huang et al., 2004; Oliet et al., 2001; Omrani et al., 2009; Rothstein et al., 1996; Takayasu et al., 2006; Tanaka et al., 1997; Tsukada et al., 2005; Tzingounis and Wadiche, 2007). Given our finding that decreased GLT1 expression levels in M1 astrocytes affected the learning and execution of movement trajectories, we examined the effects of GLT1 astrocyte deficiency on M1 layer 2/3 neuron activity *in vivo*. Previous studies have shown that in WT mice, layer 2/3 neurons show plasticity associated with learning the lever push task, with the emergence of an ensemble of correlated neurons associated with the learned movement (Peters et al., 2014). We used two-photon imaging and the calcium indicator GCaMP6s to record the calcium activity of M1 layer 2/3 neurons during the lever push task in expert animals (Figure 6A,B**, Supplemental Video 2)**. We found that the average neuronal activity pattern during successful trials was similar in WT and GLT1 mice, with very low activity at baseline (no movement) and an elevation of the calcium signal during the lever push movement period (Figure 6B,C). In WT mice, around 20% of neurons were active on average during the movement period of each trial (Figure 6D). GLT1 mice showed an increased percentage of neurons that were active during the movement period (WT: 19.89% ± 1.89, n= 6 mice; GLT1: 29.62% ± 3.01, n= 5 mice) (Figure 6D, Supplemental Figure 6A). Neuron-to-neuron signal correlation, measured by averaging the distance correlation coefficient between the concatenated trial activity vectors of pairs of single neurons, was high for a subset of WT neurons (Figure 6E-F, Supplemental Figure 6B). This group of highly correlated neurons was not found in the GLT1 trained mice (Figure 6E-F, Supplemental Figure 6B). Thus, M1 neurons in trained GLT1 mice showed significantly reduced neuronal signal correlations compared to WT mice (WT: 0.4091 ± 0.01415, n=6 mice; GLT1: 0.3516 ± 0.01835, n= 5 mice) (Figure 6E-F).

**Figure 6:**
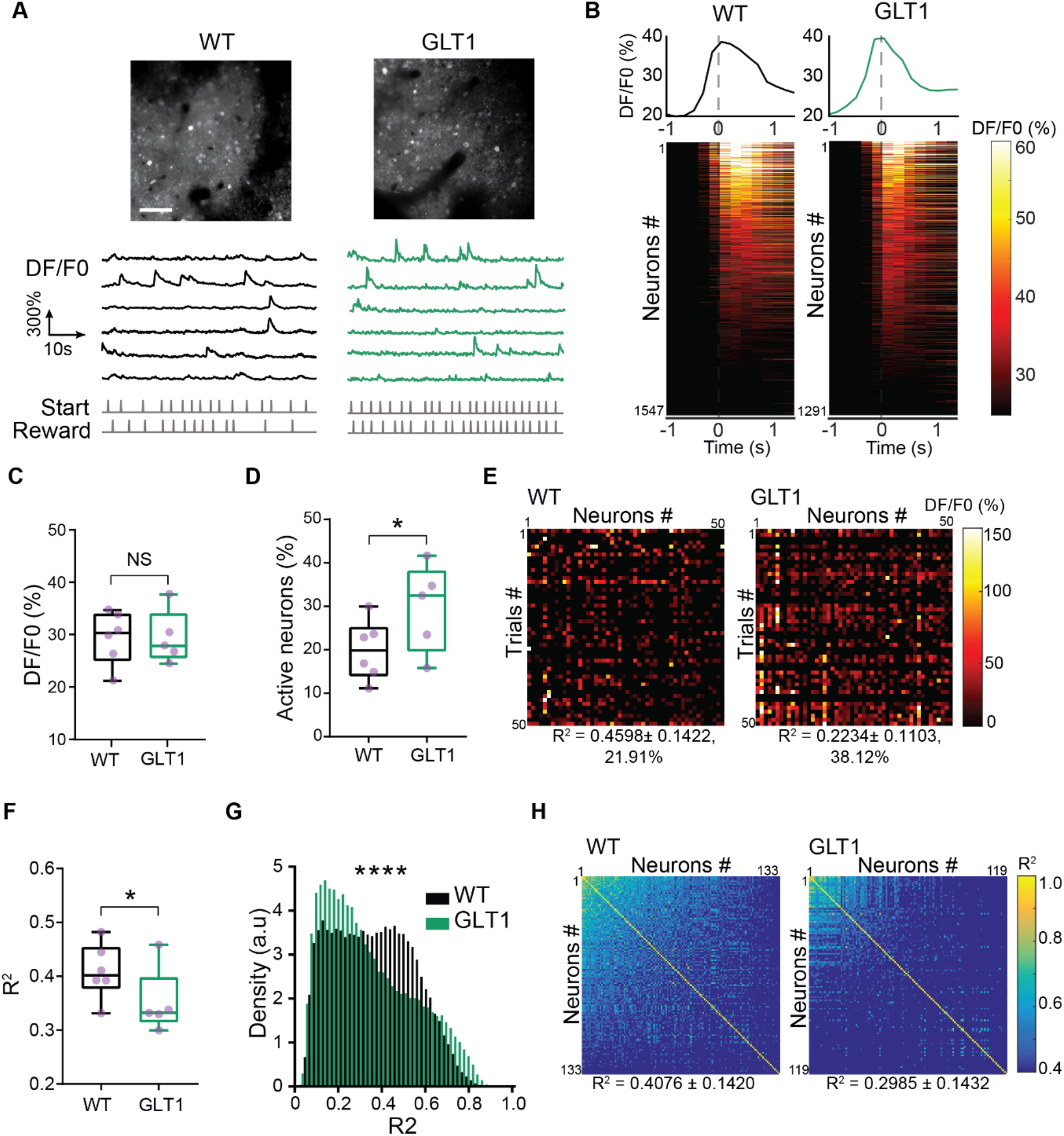
Decreased GLT1 Levels in M1 Astrocytes Reduce Neuronal Signal Correlations. **A-C.** Decreased GLT1 did not significantly change average neuronal activity. **A. Top:** Example field-of-view of neuronal GCaMP6s two-photon imaging *in vivo*. Scale bar, 25μm. **Bottom:** Example raw DF/F0 traces. **B.** Aligned trial-averaged responses of M1 layer 2/3 neurons. WT: n=1547 neurons from 15 non-overlapping fields of view from 6 mice, GLT1: n=1291 neurons from 13 non-overlapping fields of view from 5 mice, from expert session training days 10-14. **Top:** average DF/F0 trace over movement epoch. **Bottom:** normalized DF/F0 colormap; neurons are sorted by maximum activity. Zero (0) on x axis, and vertical dashed line indicate time when lever position reached the reward threshold (1mm). **C.** Average trial activity (DF/F0) (WT: mean= 29.41 ± 2.053, GLT1: mean= 29.41 ± 2.281, NS: p=0.9987, unpaired t-test). **D.** Decreased astrocyte GLT1 increased the proportion of active neurons during the movement period. Neurons were defined as active during the movement period if the activity during movement (1s period) was two standard deviations above the activity during ITI (1s period). Percentage of movement related neurons was calculated for each trial and then averaged across all trials. Percentage of active neurons during lever push was higher in GLT1 mice than WT (WT: mean= 19.89 ± 1.89, GLT1: mean= 29.62 ± 3.01, *: p=0.0103, unpaired t-test). **E.** Example colormaps of trial average activity for the first 50 trials for 50 neurons recorded in one expert training session. Neuron-to-neuron average pairwise correlation R^2^ values and percentage of active neurons are indicated below each matrix. **F-H.** Decreased astrocyte GLT1 reduced neuronal signal correlations. **F.** Trial-to-trial activity similarity was measured by the average pairwise correlation of single neuron activity vectors of concatenated trials. GLT1 mice had significantly lower average pairwise signal correlation (WT: mean= 0.4091 ± 0.01415, GLT1: mean= 0.3516 ± 0.01835, *: p=0.0204, unpaired t-test)**. G.** Density histograms of pairwise neuronal correlation distribution (****: p<0.001, Kolmogorov-Smirnov test). **H.** Example sorted correlation matrices of neuron-to-neuron average pairwise correlations for all neurons of one example session/ field of view. Neuron-to-neuron average pairwise correlation R^2^ values of the examples are indicated. For all panels, N=6 WT, 5 GLT1 mice; box plots as described in Fig. 2B.

### Gq Pathway Activation in M1 Astrocytes Increases Neuronal Signal Correlations

We also studied the calcium activity of M1 layer 2/3 neurons during movement execution in control and Gq mice at expert time points, with CNO intraperitoneal injections (Figure 7A, B). The activity patterns of the neuronal populations were similar in the two groups (Figure 7B, C). The fraction of active neurons in Gq mice injected with CNO was not significantly different from control mice injected with CNO (Figure 7D, Supplemental Figure 6C). Contrary to what we observed in the GLT1 mice, Gq mice showed increased neuron-to-neuron signal correlations, with a larger fraction of the neurons being highly correlated (CTRL+CNO: 0.323±0.0.02267 n=9, Gq+CNO: mean=0.4074±0.02154, n=6) (Figure 7E-F, Supplemental Figure 6D). Thus, Gq pathway activation in M1 astrocytes is sufficient to trigger increased correlated activity of M1 neurons.

**Figure 7:**
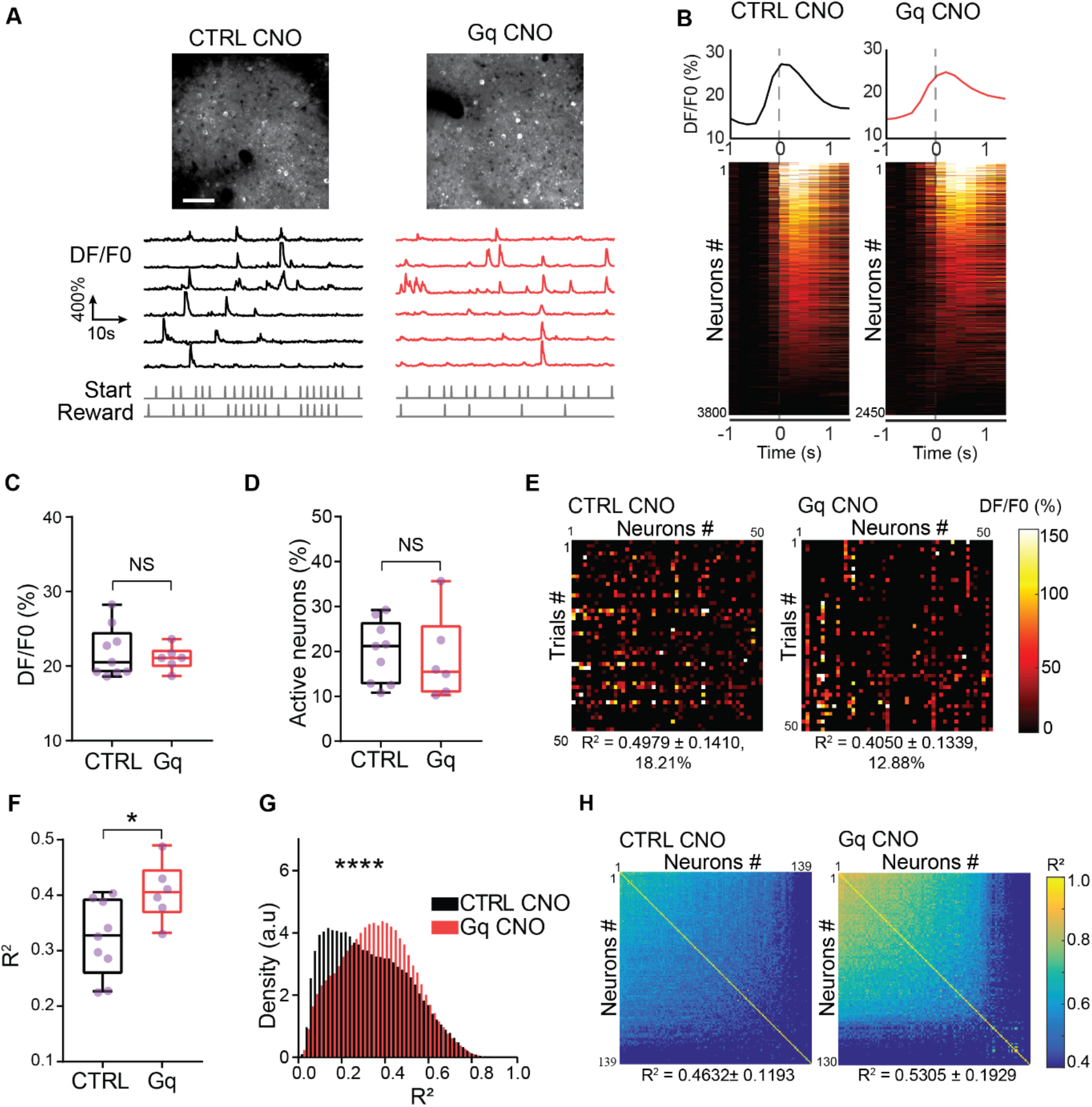
Gq Pathway Activation in M1 Astrocytes Increases Neuronal Signal Correlations. **A-C.** Astrocyte Gq activation did not significantly change average neuronal activity. **A. Top:** Example field-of-view of neuronal GCaMP6s two-photon imaging *in vivo*. Scale bar, 25μm. **Bottom:** Example raw DF/F0 traces. **B.** Aligned trial-averaged responses of M1 layer 2/3 neurons. N=3800 neurons from 9 CTRL mice injected with CNO, n=2450 neurons from 6 Gq mice injected with CNO; data from expert sessions. **Top:** average DF/F0 trace over movement epoch. Zero (0) on x axis, and vertical dashed line indicate time when lever position reached the reward threshold (1mm). **Bottom:** normalized DF/F0 colormap; neurons are sorted by maximum activity. **C.** Average trial activity (DF/F0) (CTRL+CNO: mean= 21.87 ± 1.136, Gq+CNO: mean= 21.08 ± 0.6631, NS: p=0.6077, unpaired t-test). **D.** Neurons were defined as active during the movement period if the activity during movement (1s period) was two standard deviations above the activity during ITI (1s period) and percentage of movement related neurons was calculated for each trial and then averaged across all trials. Percentage of active neurons during lever push movement was not significantly different between CTRL and Gq mice (CTRL+CNO: mean= 19.88 ± 2.286, Gq+CNO: mean= 18.41 ± 3.883, NS: p=0.7336, unpaired t-test). **E.** Example colormaps of trial average activity for the first 50 trials for 50 neurons recorded in one expert training session. Neuron-to-neuron average pairwise correlation R^2^ values and percentage of active neurons of the examples are indicated below each matrix. **F-H.** Astrocyte Gq activation increased neuronal signal correlations **F.** Trial-to-trial activity similarity was measured by the average pairwise correlation of single neuron activity vectors of concatenated trials. (CTRL+CNO: mean= 0.323 ± 0.02267, Gq+CNO: mean= 0.4074 ± 0.02154, *: p=0.0237, unpaired t-test). **G.** Density histograms of pairwise neuronal correlation distribution (****: p<0.0001; Kolmogorov-Smirnov test). **H**. Example sorted correlation matrices of neuron-to-neuron average pairwise correlations for all neurons of one example session/ field of view. Neuron-to-neuron average pairwise correlation R2 values of the examples are indicated. For all panels, n=9 CTRL+CNO, 6 Gq+CNO mice, injected intraperitoneally 30 min before all training session with low dose of CNO. Box plots as described in Fig. 2B.

### Astrocyte Manipulations Modulate M1 Neuronal Encoding of Task Parameters

Our behavioral findings showed that both astrocyte manipulations led to deficits in movement trajectory and, in the case of Gq mice but not GLT1 mice, affected hit rate and response time (Figures 4 and 5). To determine the deficit associated with these astrocyte manipulations at the neuronal coding level, we fitted decoding models of M1 neuron population activity to the push trajectory (Supplemental Figure 5; see Methods). The control groups of the GLT1 inhibition and Gq activation cohorts had similar task performances, and therefore were pooled together as the “WT” group for the decoding and encoding analyses. A support vector regression (SVR) model was used to predict the push trajectory during each training session from neuronal population spiking rate (Supplemental Figure 6A, B). For each neuronal population sample, the predictive power of the decoding model was evaluated by calculating the mutual information (M.I.) between predicted trajectory and the actual push trajectory (Supplemental Figure 6C). In WT mice, the models produced more accurate predictions of lever movement trajectories (Supplemental Figure 6C). In both Gq and GLT1 neuronal populations, the M.I. values between predicted and actual trajectories were significantly lower than that of WT neuron populations (Supplemental Figure 6C; median values: WT: 0.104, GLT1: 0.066, Gq: 0.072). These results thus indicate that in M1, astrocyte specific manipulations of glutamate transport and Gq signaling reduce neuronal population encoding of movement trajectory.

Because M1 neurons have been suggested to encode more than just directed movement signals (Doron and Brecht, 2015), we evaluated the encoding of specific behavioral features by single M1 neurons in WT, GLT1 and Gq mice. We created GLM models to predict individual neuronal activity during each trial from these specific behavioral features (Engelhard et al., 2019), and compared the prediction performance of the models between the three groups (Figure 8, Supplemental Figure 7; see Methods). The models used seven behavioral features as predictors, including two event variables - start and reward (or movement threshold); two whole trial variables - hit/miss and response time; and three continuous variables - movement trajectory, movement speed, and a step function (“moving”) indicating whether the animal started moving in a trial (Figure 8A). In WT mice, push speed, trial success (hit/miss), and response time were all predictive of the neuronal activity (median R^2^ > 1%), while the two event variables, the motion indicator and the raw movement trajectory were not very predictive (median R^2^ < 1%) (Figure 8B, **Supplemental Figure 8**). The full model with all 7 behavioral features predicted less single neuron activity variation and less encoding power for both GLT1 and Gq mice compare to WT (Figure 8C; medians: WT: 0.104, GLT1: 0.066, Gq: 0.072). Moreover, the relative contribution of different behavioral features was altered in neurons from GLT1 mice compared to WT, in favor of a proportionately increased encoding of the response time (Figure 8D). This is consistent with the impaired movement trajectory but preserved response time and success rate observed for GLT1 mice (Figure 4). In contrast, neurons from Gq mice showed an overall reduced but largely conserved relative contribution of different behavioral features (Figure 8D), suggesting a generalized reduction of the encoding of task parameters. This is consistent with Gq mice showing behavioral impairments in both task performance and movement trajectory (Figure 5).

**Figure 8:**
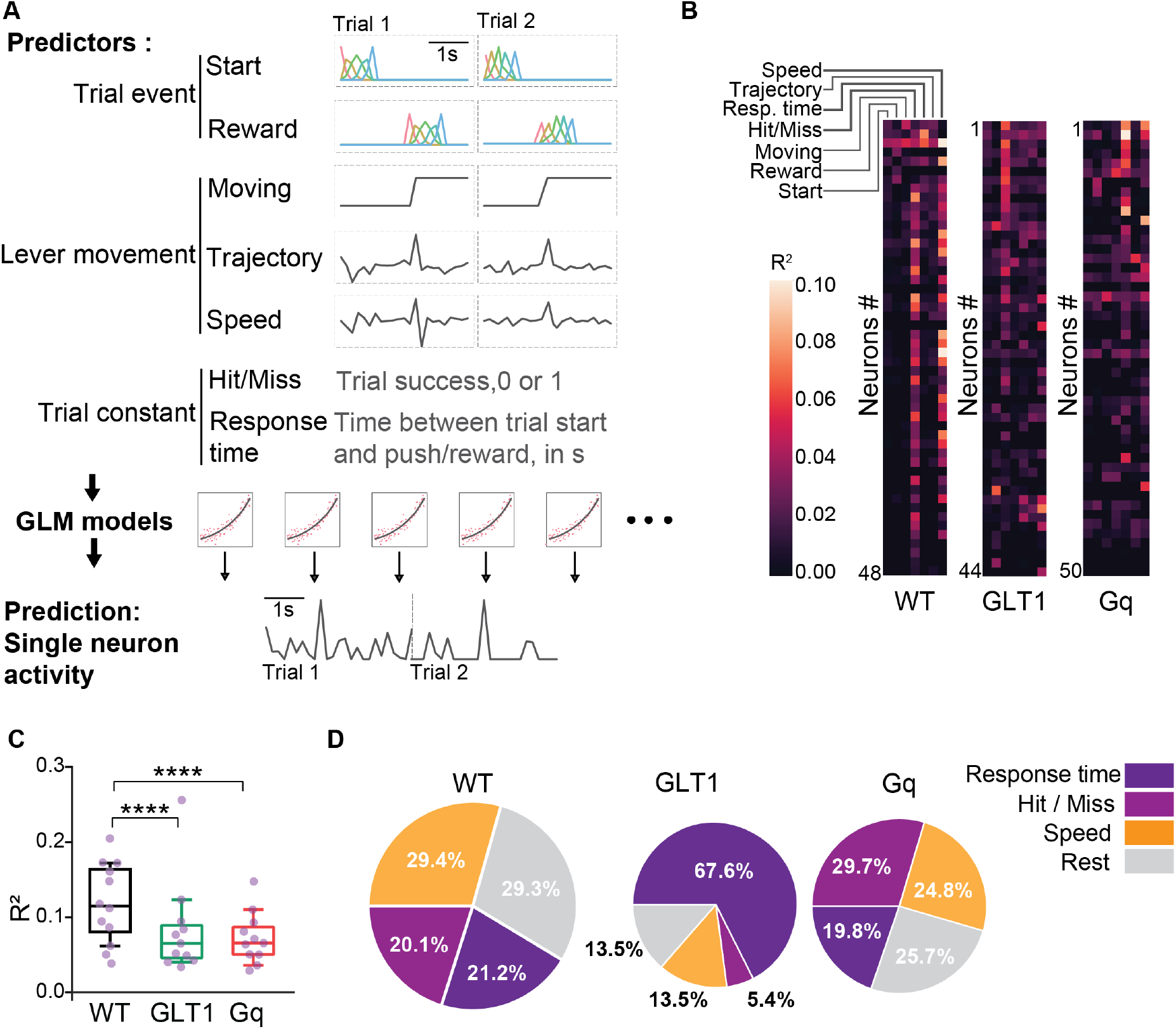
Astrocyte Manipulations Modulate M1 Neuronal Encoding of Task Parameters. **A.** The Generalized Linear Model (GLM) used to model neuronal encoding of task parameters. Predictors and predicted neuronal activity in an example trial. Seven predictors spanning trial event, lever movement, and trial constant were used. See Methods for details. One GLM model was fit for each neuron and the R^2^ between predicted and actual neuronal activity was calculated for either a full model using all the above behavioral measures as predictors, or a model with all but one behavioral feature. The difference between predictions from these two models was used to measure the contribution from the particular behavioral feature. N= 580/586/565 neurons, from 13/11/16 non-overlapping fields of view, from 5/3/5 WT/GLT1/Gq mice respectively. **B.** Representative single neurons’ encoding R^2^ values from the seven predictors for WT, GLT1 and Gq mice. Rows represent individual neurons; columns represent the contribution of individual features. **C.** Single neuron predictive power R^2^ values from all features. Both astrocyte manipulations reduced the predictive power of the full model (N=13/11/16 for WT/GLT1/Gq respectively, WT: median=0.104, GLT1: median=0.066, p = 2.15E-18, Mann-Whitney U test; WT: median=0.104, Gq: median=0.072, p = 3.14E-11, Mann-Whitney U test). Box plots as defined in Fig. 2B. **D.** Pie chart comparison of mean R^2^ across all neurons, for the most predictive task features (see Supplemental Figure 7). The size of each pie represents the mean neuronal encoding power for WT, GLT1 and Gq mice (from C). GLT1 mice showed a change in the predictors’ contribution profile compared to WT mice, with a larger relative encoding contribution of response time and relative decrease in encoding of all other features. Gq mice showed a small relative increase in neuronal encoding of trial outcome (Hit/Miss) compared to WT mice, and in general a reduced but globally conserved predictor contribution profile, suggesting a global reduction of encoding of all task parameters.

## DISCUSSION

Motor cortex is crucial for motor learning, accurate motor control and motor dexterity (Dombeck et al., 2009; Gloor et al., 2015; Harrison et al., 2012; Kawai et al., 2015; Nudo et al., 1996; Peters et al., 2017, 2014; Tennant et al., 2011). In recent years, astrocytes have emerged as key contributors to neuronal activity and plasticity (Ackerman et al., 2021; Adamsky et al., 2018; Araque et al., 1999; Corkrum et al., 2020; Haydon, 2001; Hennes et al., 2020; Kol et al., 2020; Lines et al., 2020; Mederos et al., 2019; Nagai et al., 2019; Oliveira et al., 2015; Paukert et al., 2014; Perea et al., 2014a; Poskanzer and Molofsky, 2018; Poskanzer and Yuste, 2016; Ribot et al., 2021; Santello et al., 2019; Sasaki et al., 2014; Yu et al., 2018); however, the role of astrocytes in motor cortex microcircuits *in vivo* has not been investigated so far. We show that astrocyte-specific manipulations of M1 *in vivo*, targeting glutamate clearance and Gq signaling, impacts learning and performance in a lever push task by modulation of population neuronal activity, in particular their inter-neuronal correlations and trajectory encoding, and the encoding of task parameters by single neurons. Mice expressing decreased levels of astrocyte glutamate transporter GLT1 in M1 showed normal success rate and response timing but impaired learning and execution of a stereotyped (reliable) and precise (smooth) movement trajectory. M1 neuronal population activity was strongly decorrelated and their encoding of movement trajectory was impaired. Encoding of task parameters by M1 neurons revealed a proportionately greater representation of response time, consistent with behavioral preservation of response time and success rates. Mice with astrocyte Gq signaling activation in M1 that were trained in the same task showed decreased success rate, delayed response time and impaired learning and execution of the stereotyped movement. Their altered task performance was accompanied by high levels of non-encoding M1 neuronal signal correlation, reduced population encoding of movement trajectory, and non-specific reduction of encoding of task parameters by single neurons. Using M1 as a test bed, these findings thus provide quantitative evidence for the role of astrocytes in influencing the coding of information by single neurons and neuronal populations during learning.

Our manipulations were motivated by changes in gene expression and calcium activity of M1 astrocytes during the lever push task. We observed changes in expression for a small number of individual genes and for a larger number of gene sets. Combined with the finding that calcium events in astrocytes become more coincident with the lever push movement, our results suggested that astrocytes display plasticity at the gene expression and functional levels that are associated with motor learning. Therefore, astrocyte-specific manipulations would be expected to alter behavioral and neuronal function during learning. In particular, glutamate transport stood out from the identified enriched gene sets, supporting the hypothesis that astrocyte glutamate transport has a major role in M1 during motor learning.

The astrocytic glutamate transporter GLT1 is the major glutamate transporter in the cerebral cortex. Its role in regulating glutamate availability and accumulation of extracellular glutamate has been well documented, along with its role in limiting glutamate spillover to neighboring synapses and extrasynaptic receptors (Arnth-Jensen et al., 2002; Asztely et al., 1997; Bergles et al., 1999; Diamond and Jahr, 1997; Rothstein et al., 1996; Tanaka et al., 1997). Increasing evidence also links GLT1 to localized calcium events. While several pathways potentially contribute to such events, including membrane channels (Rungta et al., 2016; Shigetomi et al., 2013), metabotropic and ionotropic receptor activation (Agulhon et al., 2012), and membrane transporters (Bernardinelli et al., 2006), recent evidence has demonstrated that mitochondrial calcium efflux accounts for many of the observed calcium events *in vivo* (Jackson et al., 2014; Robinson and Jackson, 2016). A working hypothesis is that synaptically released glutamate is removed via GLT1 (with a net influx of sodium ions) and converted to glutamine, metabolized to generate ATP by local mitochondria, which generates focal mitochondrial calcium signals (Griffiths and Rutter, 2009; Jackson and Robinson, 2015; Stephen et al., 2015). Consistent with these various roles of GLT1, we observed that decreasing GLT1 levels in M1 layer 2/3 astrocytes triggered an increase in the proportion of active neurons during the movement epoch of the task. Moreover, the neuronal populations failed to form an ensemble of highly correlated neurons, which has previously been associated with motor learning (Peters et al., 2014). GLT1 mice failed to learn a stereotyped and precise lever push movement but showed preserved hit/miss performance in the task. This phenotype is similar to the motor learning deficit observed after pre-learning M1 lesions in rodents (Kawai et al., 2015; Peters et al., 2014). We also show here that GLT1 reduction triggers complex changes in neuronal activity, including not only increased proportions of active neurons with reduced inter-neuronal correlations but also reduced population encoding of movement trajectory and altered single neuron encoding of task parameters.

In contrast, astrocyte-specific Gq signaling activation in M1 astrocytes triggered an increase in neuronal signal correlation that appeared to be non-informative. This suggests a crucial role for astrocytes in decorrelating neurons through Gq-dependent mechanisms. The behavioral phenotype was accompanied by a significant increase in response delay, decrease in the fraction of successful trials (hit rate) and decrease in stereotypy of the push trajectory. The failure of Gq-activated astrocytes to decorrelate neuronal activity in M1 layer 2/3 during motor learning may affect downstream neurons in charge of task execution, leading to delayed responses and reduced task performance. The behavioral phenotype was rapidly improved when astrocyte-specific Gq activation was stopped, suggesting that the perturbation was transient and reversible, and affected mechanisms of execution during motor learning rather than learning *per se*. The response time was the task parameter that showed the largest change following Gq activation of M1 astrocytes, and was greatly improved but not totally restored in the CNO withdrawal group. One hypothesis would be that the Gq-DREADD construct by itself (without CNO) had an effect. We thus performed a control experiment with Gq-DREADD mice injected with saline throughout the learning of the task, and did not see any difference from the control group. Another factor could be the existence of a lasting effect of the CNO despite withdrawal, either by direct residual presence in the cortex or by indirect effect on task performance through lasting functional or structural cellular changes. Gq-GPCR is known to trigger intracellular calcium elevation through IP3-induced calcium release from the ER (Agulhon et al., 2013; Clapham, 2007; Mizuno and Itoh, 2009). We found that M1 astrocyte Gq activation was associated *in vivo* with an increase in intracellular calcium, likely to trigger a saturation of calcium signals and consequently a decrease in frequency of calcium events. This result is consistent with a recent study demonstrating by similar methods a decrease in calcium dynamics in Gq activated cortical astrocytes (Vaidyanathan et al., 2021). We note that although Gq-DREADD is currently one of the most relevant tools available to study astrocyte Gq pathway activation and to modulate astrocyte function; how accurately it reflects Gq pathway activation physiologically *in vivo* remains to be determined.

It was demonstrated that calcium signaling inhibition in hippocampal astrocytes prevented the diversity of neuronal presynaptic strengths (Letellier et al., 2016). Moreover, a study showed that reduction of astrocyte calcium signals in the striatum greatly increased the inter-neuronal correlation of striatal medium spiny neurons during non-grooming episodes (Yu et al., 2018). Our finding that astrocytic Gq activation and the associated reduction of astrocyte calcium dynamics increase non-encoding neuronal correlations is consistent with these studies. Together, they support a role for astrocytes in the maintenance of neuronal decorrelation and synaptic strength heterogeneity.

Given our finding that modifications of astrocyte gene expression and calcium events are associated with motor learning, one possibility is that astrocyte manipulations may disrupt astrocyte-neuron plasticity during learning. In support of this idea, we observed that decrease in astrocyte GLT1 levels and activation of astrocyte Gq signaling both prevented a number of gene expression changes during motor learning. Our study also revealed that activation of astrocyte Gq signaling triggered an increase in *Slc1a2*/GLT1 expression. Gq activation thus may be expected to have contrasting effects compared to GLT1 inhibition in M1 astrocytes. While some effects were symmetrical, others were not, indicating that balanced astrocyte-neuron activity is critical for proper operation of brain circuits.

## MATERIAL & METHODS

### Experimental Model

All experimental procedures performed on mice were approved by the Massachusetts Institute of Technology Animal Care and Use Committee, and conformed to National Institutes of Health guidelines for the Care and Use of Laboratory Animals. Adult mice (2 to 4 months old, C57BL/6J background) were housed on 12-hour light/dark cycle, group housed before surgery and singly housed afterwards. Male and female mice were used. The following mouse lines were used: C57BL/6J wild-type (JAX Stock #000664, Jackson Laboratory, Bar Harbor, ME), CaMKII;mTTA;GCAMP6s (mTTA;GCAMP6s: Ai94(TITL-GCaMP6s)-D;ROSA26-ZtTA, JAX Stock #024112, and CaMKII-cre: B6.Cg-Tg(Camk2a-cre)T29-1Stl/J, JAX Stock #005359), GFAP;GCaMP5G (GFAP-cre: B6.Cg-Tg(Gfap-cre)77.6Mvs/2J, JAX Stock #024098 and GCaMP5G, Polr2atm1(CAG-GCaMP5g,-tdTomato)Tvrd, JAX Stock #024477), Aldh1l1;GCaMP6f-Lck (Aldh1l1-cre: B6;FVB-Tg(Aldh1l1-cre)JD1884Htz/J, JAX Stock #023748, and GCaMP6f-Lck: C57BL/6N-Gt(ROSA)26Sortm1(CAG-GCaMP6f)Khakh/J, JAX Stock #029626). The GLT-1 flox line (Cui et al., 2014) was a gift from Kohichi Tanaka.

### Stereotactic virus injection and craniotomy

Surgeries were performed aseptically, under isofluorane anesthesia while maintaining body temperature at 37.5C. Mice were given preemptive analgesia (slow release buprenex, subcutaneous, 0.1mg/kg). Scalp hairs were removed with hair-remover cream, skin was sterilized with 70% ethanol and betadine, and portion of the scalp was removed. Mice were head-fixed in a stereotaxic frame (51725D, Stoelting Co., Wood Dale, IL). A 3mm diameter round craniotomy was performed over the left motor cortex (0.3mm anterior and 1.5mm lateral to bregma) and a 200nL volume of virus solution (titer of 10-12 virus molecules per ml) was injected 300μm below the pial surface at 50nL/min with a thin glass pipette and a stereotaxic injector (QSI 53311, Stoelting). Following each injection, the glass pipette was left in place for 15 additional minutes and was then slowly withdrawn to avoid virus backflow. The following viruses were used: AAV8-GFAP-hM3D(Gq)-mCherry (UNC Vector Core), AAV5.GFAP.Cre.WPRE.hGH (Penn Vector Core, AV-5-PV2408), AAV1.Syn.GCaMP6s.WPRE.SV40 (Penn Vector Core, AV-1-PV2824). Finally, a cranial window made of 3 round coverglasses (1x 5mm diameter CS-5R, and 2x 3mm diameter CS-3R, Warner Instruments, Hamden, CT) glued together with UV-cured adhesive (NOA 61, Norland, Jamesburg, NJ) was implanted over the craniotomy and sealed with dental cement (C&B Metabond, Parkell, Brentwood, NY). For head fixation during the behavioral task and/or calcium imaging, a headplate was also affixed to the skull using dental cement (C&B Metabond, Parkell). Postoperative analgesic was provided (Meloxicam, subcutaneous, 1mg/kg) and recovery was monitored for a minimum of 72 hours after surgery. Animals were allowed to recover for at least five days before starting the water restriction for behavioral experiments. Upon completion of experiments, we verified that targeting of the motor cortex region was successful by immunohistochemical techniques and fluorescence confocal imaging. Animals for which viral delivery was mis-targeted or failed were excluded.

### Behavioral testing

Water restricted mice were head fixed and trained daily on the lever push task (Peters et al., 2014), modified as follows. The lever was built using a piezoelectric flexible force transducer (LCL-113G, Omega Engineering, Norwalk, CT) attached to a brass rod and could be reached easily by mice using their right paw. Another fixed brass rod was placed in front of the left paw. The voltage from the force transducer, which is proportional to the lever position, was continuously recorded. Lever press was defined as crossing of a 1mm threshold. A tone marked the beginning of a trial, with a 5 sec response period. A lever press past the threshold triggered a 6μL water reward and the start of a 2.62s reward time followed by an inter-trial interval (ITI). Failure to press during the 5s response period triggered a loud white noise and a 2.62s timeout period followed by the ITI. Lever presses during the ITI were punished by delaying the start of the next trial until a full second of time passed without any lever movement. The system was controlled by MATLAB (MathWorks, Natick, MA) using the Psych-Toolbox.

### CNO administration

CNO (Enzo Life Sciences, Farmingdale, NY) was dissolved in saline injectable sterile solution (0.9% sodium chloride) at a low 0.1mg/kg concentration. The CNO solution or saline control was intraperitoneally injected 30 min before each training session. The CNO concentration used was very low compared to published studies and did not induce any seizure.

### Two-photon microscopy

Mice were head fixed and GCaMP fluorescence imaging of the left motor cortex (0.3mm anterior and 1.5mm lateral to bregma) was performed through the cranial window, 2-6 weeks post virus injection and after 3 days of habituation (consisting in daily 10min passive sessions). A Prairie Ultima IV two-photon microscopy system was used with a galvo-galvo scanning module (Bruker, Billerica, MA). 910nm wavelength excitation light was provided by a tunable Ti:Sapphire laser (Mai-Tai eHP, Spectra-Physics, Milpitas, CA) with dispersion compensation (DeepSee, Spectra-Physics). For collection, GaAsP photomultiplier tubes (Hamamatsu, Bridgewater, NJ) were used. Images were acquired using PrairieView acquisition software.

*GCaMP6f-lck:* to detect all astrocyte calcium events including smaller and faster ones, we used a 25x/1.05 NA microscope objective (Nikon) combined with 4x optical zoom and acquired image sequences at 11Hz for 10min during each novice or expert training session of awake mice performing the lever push task. Three 300×150 pixel (80.3 x 40.2 µm) rectangular fovs were imaged per 10min session, with each fov imaged for 200s (1/3 of the session) sequentially.

*GCaMP6s:* A 16x/0.8 NA microscope objective (Nikon, Tokyo, Japan) was combined with 2x optical zoom to achieve simultaneous imaging of a large number of neuronal somas, and image sequences were acquired at 5Hz. A 521×236 pixel (274 x 274 µm) square field of view (fov) was imaged for 10min during each expert training session of awake mice performing the lever push task.

*GCaMP5G:* A 16x/0.8 NA microscope objective (Nikon) was combined with 2x optical zoom to achieve simultaneous imaging of a large number of astrocytes, and image sequences were acquired at 5Hz. A 521×236 pixel (274 x 274 µm) square fov was imaged for each 10min passive imaging session in awake untrained mice.

### Astrocyte activity image analysis

Astrocyte calcium activity during novice or expert training sessions of awake mice performing the lever push task was analyzed in ALDH1L1; GCaMP6f-lck mice as follow.

#### Motion correction

After acquisition, time-lapse imaging sequences were corrected for x and y movement using the template-matching NoRMCorre algorithm(Pnevmatikakis and Giovannucci, 2017).

#### Event detection

Spatiotemporal events were detected using the AQuA algorithm (Wang et al., 2019). Briefly: fluorescence signals 2 standard deviations above baseline (F0) and at least 16 pixels (= 1.145 µm^2) in size were identified. Foreground signals in neighboring pixels in the spatiotemporal directions (XYZ) were then grouped into an event based on similarity in onset time, offset time, and proximity. The total pixels in the XYZ planes that are grouped together are considered as an event and are used to calculate event features. Z-scores for each event were calculated from the average of all normalized pixel values in the event and events with Z-scores less than 3 were considered as noise and excluded from the analysis.

#### Number of events

For each training session, XYZ events were detected as described above from each fov and the total number of events detected in the three 200s sequences was summed to yield the total number of events for the 10min imaging/behavior session.

#### Event area

The area of the event was calculated by measuring the area of the spatial footprint of the XYZ event in the XY direction.

#### Event amplitude

The amplitude of an XYZ event was calculated as the maximum detected DF/F0 for the XY event within the event time period (i.e., the maximum across the Z dimension), with DF/F0 = 100*(F – F0)/F0, where F is the average XY signal and F0 is the baseline fluorescence value for the entire video.

#### Percentage of trials with coincident events

Trials with coincident events were detected first by identifying the closest event that occurred after the lever push, and then determining whether one or more events also occurred during this movement-associated event.

#### Trial-averaged activity

A single DF/F0 vector for the entire imaging session was generated for each event using the AQuA package. Briefly, this consisted in removing contributions from other events, which were then imputed with nearby values. The trial-averaged activity during movement for each successful trial was defined as the average DF/F0 value across all events during a 2s epoch starting 1s before movement onset.

Changes in astrocyte calcium induced by Gq pathway activation were analyzed in GFAP; GCaMP5G mice as follow. After acquisition, time-lapse imaging sequences were corrected for x and y movement using the template-matching ImageJ plugins. Regions of interests (ROI) were automatically identified using CaSCaDe (Agarwal et al., 2017). The baseline fluorescence F0 was calculated as the 25^th^ percentile. DF/F0 (= 100*(F – F0)/F0) was calculated where F is the ROI average fluorescence and F0 the baseline fluorescence. DF/F0 peaks with values three standard deviations above the average DF/F0 were considered as calcium elevation events. The event amplitude was defined as its maximum DF/F0 value.

### Neuronal activity image analysis

GCaMP6s fluorescence from the upper layers of the left motor cortex was acquired as described above in CaMKII;mTTA;GCAMP6s transgenic mice or alternatively, AAV1.Syn.GCaMP6s.WPRE.SV40 injected wildtype mice. We used GCaMP6s for this study due to its higher SNR that can better capture the motion information encoded in M1 neurons, and especially, compared to GCaMP6f, its larger response amplitude, lower variability and thus greater single-spike detectability when used to infer spikes (Huang et al., 2019; Wei et al., 2020). After acquisition, time-lapse imaging sequences were corrected for x and y movement using template-matching ImageJ plugins. Regions of interests (ROI) were automatically identified using Suite2P (Pachitariu et al., 2016) and then manually curated. Alternatively, neuronal ROIs were manually selected. The fluorescence intensity in time for each ROI was then averaged. The DF/F0 (= 100*(F – F0)/F0) was calculated where F is the average signal and F0 the mode of the signal.

#### Average activity

Activity during movement or during ITI for each successful trial was defined respectively as the average DF/F0 across a 1s epoch starting at movement onset or as the average DF/F0 across the final 1s of the ITI (no movement). The “average activity” of a neuron was calculated as the average across all the successful trials of the activity during movement as defined above.

#### Movement related active neurons

For each trial, a neuron was considered “active” if the maximum DF/F0 during movement was two standard deviations above the average DF/F0 during ITI. Percentage of movement related neurons was calculated for each trial and then averaged across all trials.

#### Neuron-to-neuron correlation

Neurons active for more than 10% of the trials were included in the analysis. For each neuron, activity data vectors during movement for all trials were concatenated into one vector. The pairwise distance correlation coefficient of two vectors was then computed to estimate neuron-to-neuron correlation.

#### Behavioral encoding model

We used encoding models to test and compare the accuracy of using different behavior variables to predict the variability in the neuronal activity during the lever push task trials in fully trained mice. We employed a Generalized Linear Model (GLM) as described in Engelhard et al., 2019 (Engelhard et al., 2019) modified as follow (see also Fig. 8A). Behavioral events, lever trajectory and neuronal activity (DF/F) during a 5s period after the start of each correct trial were used. For each training session, data vectors for all trials were concatenated into one vector before fitting the model. We extracted 7 basic features of the behavioral data in the model, and further expanded them temporally to facilitate a linear model. Three types of predictors were used: events, trial constants and continuous variables, as follows. Two events, start of trial and reward were included (reward was immediately given when the lever was pushed past the threshold). These events were converted into continuous variables with the same sampling rate as the neuronal activity, by convolving each with a 7-degrees-of-freedom regression spline basis set. Trial constants were single variables specific to a trial, including trial status (hit/miss, scored as 1 or 0 respectively) and response time as a number (time in s). These trial constants were converted to timeseries by convolving with a step function lasting the duration of the trial. Continuous variables included lever trajectory, lever speed and lever motion (“moving”), each raised to a 3^rd^ degree polynomial. A special case was lever motion, which was an in-trial step function that was set to 1 before the movement onset and 0 after onset. This predictor encoded whether or not a movement was occurring, but did not differentiate how long the movement epoch was for each trial (details of predictor transformation similar to Engelhard et al., 2019). The expanded predictors were scaled (z-scored) and fitted to a linear model for each neuron, regularized with an elastic net penalty. The accuracy of each GLM model was assessed by 5-fold splits cross-validation (80% of data for training set, 20% for testing set). The encoding power R^2^ was calculated for each prediction from the fitted model. We quantified the relative contribution of each behavioral variable to single neuron activity by determining how the performance of the encoding model was reduced (decrease of R^2^) when each variable was excluded of the predictor set of the model, the model was kept as-is while setting the weights of the excluded feature to 0. When excluding a variable, its derivative/expanded predictors were taken out as well.

#### Decoding analysis

To evaluate and compare the capacity of M1 neurons to encode the forelimb push trajectory, we tested a decoding model to predict the push trajectory from the neuronal population activity. The calcium activity (DF/F) was deconvolved with an adaptive kernel to obtain an estimate of spiking activity (Vogelstein et al., 2010). We fitted a linear decoding model to the entire duration (600s) of each training session, as well as to concatenated push trajectories alone, to predict the forelimb push trajectory from population spiking activity of the most informative 20 recorded M1 layer 2/3 neurons for each session. After evaluation of various M1 decoding models, we chose the support vector regression (SVR) model for measuring the encoding information in M1 neurons for its stability, (i.e. higher reliability with random split validation), which allowed us to compare neuron encoding capacities of populations from different animals in different groups. Specifically, a SVR model with radial basis function (RBF) kernel was used. This model has two main hyper-parameters: γ, the scaling parameter of the kernel, and C, the regularization parameter. They were optimized through grid search for the average prediction performance (as discussed below) across all cases in all treatment groups, and the same hyper-parameters were used for all cases (γ = 1E-4, C = 12). For each session, a continuous segment of time making up 10% of the entire session starting at a randomized time point (typically, 60s segment in a 600s training session) was used as the test period for model fitting, during which both behavioral and neuronal data were taken as the test dataset. The rest of the data was spliced together as the training set. To quantify the similarity between the model predicted trajectory and the actual trajectory, we used a modified version of the Non-Parametric Entropy Estimation Toolbox (NPEET) package (Ver Steeg and Galstyan, 2013). Briefly, we used a continuous estimation of mutual information by the average data point distance to the k-th neighbor (usually used with k = 3). This mutual information between predicted trajectory and actual trajectory was used as the metric of encoding capacity of M1 neuron populations.

### Immunohistochemistry

Mice were transcardially perfused with 0.9% saline followed by 4% paraformaldehyde (PFA) in PBS. Coronal sections were cut to a thickness of 50mm using a vibratome (VT1200S, Leica, Wetzlar, Germany) and incubated for 1h in blocking solution (0.1% Triton + 3% BSA in PBS), then overnight in blocking solution with the following primary antibody: 1:1000 mouse anti S-100 β subunit (S2532, Sigma-Aldrich, St. Louis, MO). Sections were washed and then incubated for 2h in blocking solution with the following secondary antibody at a 1:500 dilution: goat anti-mouse 647nm (A21235, ThermoFischer, Waltham, MA). Sections were washed in PSB, then mounted on slides in hard set mounting medium containing 4,6-diamidino-2-phenylindole (DAPI) (Vectashield, H-1500, Vector Laboratories, Burlingame, CA). A confocal system (TCS SP8, Leica) was used to image the fluorescence of GCaMPs, mCherry and S100b immunostaining, using 10×/0.40, 20×/0.75, or 63×/1.40 objectives (magnification/numerical aperture, Leica) and the LAS X Acquisition Software (Leica).

### Western Blot

Mice were deeply anesthetized under isoflurane, decapitated and their brain immediately extracted and dissected on ice-cold 0.9% saline. Cortices were dissected (∼4mm^3^ samples) and meninges removed. Left and right M1 cortex biopsies were flash frozen and stored at −80C. Frozen samples were later homogenized in ice-cold RIPA buffer (#89901, ThermoFisher) supplemented with phosphatase inhibitors (PhosSTOP, Roche, Indianapolis, IN) and protease inhibitors ( cOmplete, Mini, EDTA-free, Roche) using a high-speed homogenizer (Fast-Prep-24 5G Instrument, MP Biomedicals, Irvine, CA). BCA protein assay kit (ThermoFisher Pierce BCA Protein Assay) was used to determine the protein concentration. After denaturation at 95C for 10 min, samples were loaded on 4–15% polyacrylamide gels (BioRad Laboratories, Hercules, CA), transferred to PVDF membranes (MilliporeSigma, Burlington, MA), and immunoblotted for protein expression using the following antibodies: guinea pig anti-GLT1 at 1:25,000 (Millipore AB1783) and mouse anti-beta actin at 1:20,000 (Sigma-Aldrich A1978), and the following fluorescent secondaries: donkey anti-rabbit IRDye 800CW (LI-COR, Lincoln, NE) at 1:10,000 and Goat anti-Mouse IRDye 680RD (LI-COR). Immunoreactive bands were imaged with LI-COR Odyssey and quantified using ImageJ software. Protein levels were normalized to Actin levels. Normalized values were standardized by using the ratio of the left hemisphere, injected with viral solution, to the right, non-injected, hemisphere.

### Quantitative RTqPCR

Left M1 cortices were extracted as described above and homogenized in TRIzol (Invitrogen, Waltham, MA) using a high-speed homogenizer (MP Biomedicals Fast-Prep-24 5G Instrument). Total RNA was isolated using phenol-chloroform extraction and then purified and concentrated using ethanol precipitation and washing on a silica column (RNA Clean & Concentrator-5, Zymo Research, Irvine, CA). Total RNA samples were reverse transcribed (ThermoFisher SuperScirpt IV Vilo). The RTqPCR were perfomed using a QuantStudio™ 3 System (ThermoFisher) with SYBR Green enzyme mix (ThermoFisher PowerUp SYBR Green Master Mix). The following primers were used: Glyceraldehyde 3-phosphate dehydrogenase (*Gapdh*) Forward: AAGAGAGGCCCTATCCCAAC, Reverse: GCAGCGAACTTTATTGATGG; peptidylprolyl isomerase A (*Ppia*) Forward: GTGACTTTACACGCCATAATG, Reverse: ACAAGATGCCAGGACCTGTAT; solute carrier family 1, member 2 (*Slc1a2*) Forward: GAACGAGGCCCCTGAAGAAA, Reverse: CCTGTTCACCCATCTTCCCC; solute carrier family 1, member 3 (*Slc1a3*), Forward: GTAACCCGGAAGAACCCCTG, Reverse: GTGATGCGTTTGTCCACACC; solute carrier family 6, member 1 (*Slc6a1*), Forward: CACTCTGTTCTGGTGTCCCC, Reverse: GGGAAGCTTAATGCCAGGGT; solute carrier family 6, member 11 (*Slc6a11*), Forward: ATGATGCCCCTCTCTCCACT, Reverse: TACCACGGCTGTCACAAGAC; solute carrier family 6, member 6 (*Slc6a6*), Forward: TTCAGACAACAGACACGCGA, Reverse: CTCGGCAGCAACCAGGTC; testican-2 (*Spock2*), Forward: AGGTCACATTTCAGCCACGA, Reverse: TTGATGTCCTTCCCTCCACC, basigin (*Bsg*) Forward: GGCGGGCACCATCCAAA, Reverse: CCTTGCCACCTCTCATCCAG. Every sample was run in technical duplicate or triplicate. Relative expression was quantified using the ΔΔCp method.

### RNAseq

Wild-type mice were water restricted and then trained for 0 (naive), 3 (novice) or 19 (expert) days. To match stress levels, all three groups were water-restricted and head-fixed for the same duration as the expert mice. M1 cortices were then dissected as described above, but using ice cold ACSF (120 mM, KCL 3mM, NaHCO3 26.2mM, MgSO4 2mM, CaCl2 0.2mM, D-Glucose 11.1mM, HEPES 5mM) bubbled with oxygen and supplemented with AP5 (0.02mM) and CNQX (0.02mM) instead of PBS. Cells were dissociated using Miltenyi Biotec Neural Tissue Dissociation kit – Postnatal Neurons (130-094-802) and gentleMACS Dissociator following manufacturer protocols. Cell suspension was depleted of microglia and myelin debris (Myelin Removal Kit, 130096733, Miltenyi Biotec, Bergisch Gladbach, Germany), then astrocytes were isolated using the Miltenyi Biotec’s anti-ACSA-2 magnetic cell sorting kit and protocol (Miltenyi Biotec, 130097678). RNA was purified and concentrated with proteinase K cell digestion, ethanol precipitation and washing on a silica column (Zymo Quick-RNA FFPE). RNA concentration and quality were assessed with Agilent 2100 Bioanalyzer. Indexed cDNA libraries were generated using the SMARTer Stranded Total RNA-Seq Kit v2 (#634411, Illumina, San Diego, CA) and multiplexed sequencing was performed on Illumina HiSeq 2000. Reads were aligned to the mouse mm9 genome using the TopHat sliced read mapper (Trapnell et al., 2012). Fragment counts were obtained using the Cufflinks pipeline (Trapnell et al., 2012). Genes with fragment counts above 20 kpm were selected for further analysis. To remove unwanted variation, normalization was implemented using the Bioconductor packages EDASeq (Risso et al., 2011) and RUVSeq (Risso et al., 2014). Differential expression analysis was performed using Bioconductor package EdgeR (Robinson et al., 2009). The gene ontology (GO) analysis of DEGs was performed using PANTHER (Mi et al., 2019). The Gene Set Enrichment Analysis was performed using Bioconductor EdgeR Camera package (Wu and Smyth, n.d.). The data is available at the Gene Expression Omnibus repository (GEO: GSE156661).

### Statistical Analysis

All experiments included at least three replicates, with each replicate being a mouse-averaged value if not otherwise stated. Line plots and bar graphs show mean ± standard error of the mean (SEM). Box plot bars represents median, the box extends from the 25th to 75th percentiles, and whiskers show 10th to the 90^th^ percentile. No statistical methods were used to pre-determine sample size. We used sample sizes similar to literature in the field. Sample sizes provided at least 80% power to detect the experimental effect. For datasets with two data groups, groups were compared using Student’s two-tailed t-tests or Mann-Whitney U tests. Paired comparisons were performed with paired t-test or paired Wilcoxon rank sum test. Comparisons of cumulative distributions were performed using a nonparametric Kolmogorov-Smirnov test. For datasets with three of more data groups, groups were compared using one-way ANOVA with multiple comparisons test. Datasets with different treatment groups or different models built on grouped data were compared with (treatment/model) x astrocyte group two-way ANOVA, either with or without linear mixed model for individual animal effects, as indicated in the text. Results of statistical tests are reported in the figure legends. Values and replicate numbers are defined in the figure legends.

### Materials Availability

Further information and requests about data, resources and reagents should be directed to and will be fulfilled by Mriganka Sur.

## ACKNOWLEDGEMENTS

We thank Vaibhavi Shah, Taylor Johns, Vincent Pham, Liadan Gunter and Austin Sullins for their technical help. We thank all members of the Sur lab for their support. We thank the MIT Division of Comparative Medicine for animal care. We thank the MIT BioMicroCenter for performing the RNAseq, and the Picower Institute Bioinformatics Core Facility for analysis support. We thank Kohichi Tanaka for providing the GLT1 flox mouse line, which contributed crucially to this study. This work was supported by the NIH (grants R01EY028219, R01DA049005), MURI-ARO Proposal 78249-NS-MUR, and the Simons Foundation Autism Research Initiative through the Simons Center for the Social Brain (M.S.). C.D. was supported by awards from the Bettancourt-Schueller Foundation and Philippe Foundation, and by fellowships from the Simons Center for the Social Brain and Picower Foundation. K.L. was supported by the Rettsyndrome.org foundation’s mentored training fellowship #3213. J.S. was supported by a NIH Ruth L. Kirschstein Postdoctoral NRSA (F32NS110481).

## AUTHOR CONTRIBUTIONS

C.D. and M.S. designed the experiments. C.D. performed the behavioral experiments, stereotaxic injection and cranial window surgeries, M1 microdissections, immunohistochemistry, western blotting and two-photon calcium imaging experiments and analysis. P.G. performed behavioral experiments, astrocyte purification and RNA extractions. C.D. processed and analyzed the RNA-seq data. J.S. performed cranial window surgeries and astrocyte imaging experiments and analyzed the data. C.D. and K.L. processed and analyzed the two-photon neuronal imaging data. K.L. performed the decoding and encoding analysis. C.D., K.L. and M.S wrote the manuscript.

## SUPPLEMENTAL INFORMATION

### Supplemental Figures 1-8

**Supplemental Figure 1:**
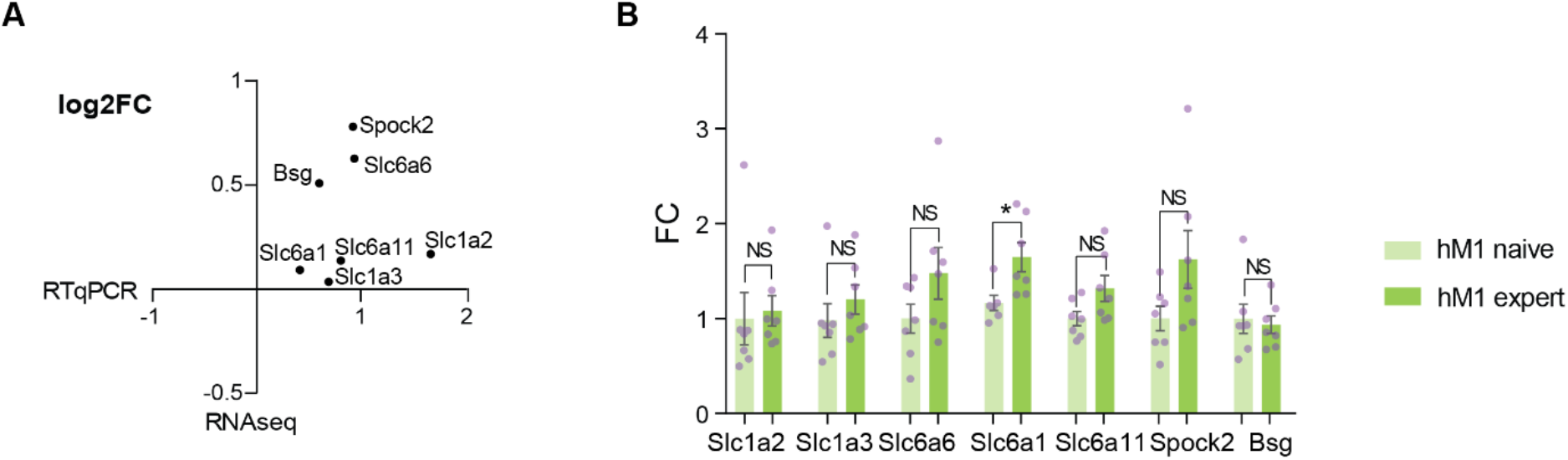
Motor Learning Leads to Modification of Gene Expression Profiles. **A.** XY plot of logarithm of fold change (log2FC) in WT expert mice relative to naïve mice, as measured in RNAseq experiment (y axis) compared to RTqPCR experiments (x axis), using independent samples. The two independent experiments showed the same trend of gene expression changes in expert mice compared to naïve mice. N=3-6 mice per group for RTqPCR, 6 mice per group for RNAseq. **B.** Bar plot (mean± SEM) representing the expression fold change (FC) of selected genes in the hindlimb motor cortex (hM1) of WT naïve and expert mice (n=7 mice per group). Slc1a2 (naïve 1±0.2766, expert 1.082±0.1600, NS: p=0.8018, unpaired t-test), Slc1a3 (naïve 1±0.1777, expert 1.201±0.1527, NS: p=0.3632, unpaired t-test), Slc6a6 (naïve 1±0.1516, expert 1.478±0.2721, NS: p=0.1511, unpaired t-test), Slc6a1 (naïve 1±0.0802, expert 1.648±0.1521, *: p=0.0217, unpaired t-test), Slc6a11 (naïve 1±0.07358, expert 1.319±0.1362, NS: p=0.0620, unpaired t-test), Spock2 (naïve 1±0.1275, expert 1.624±0.3058, NS: p=0.084, unpaired t-test), Bsg (naïve 1±0.1541, expert 0.9357±0.091, NS: p=0.7316, unpaired t-test).

**Supplemental Figure 2:**
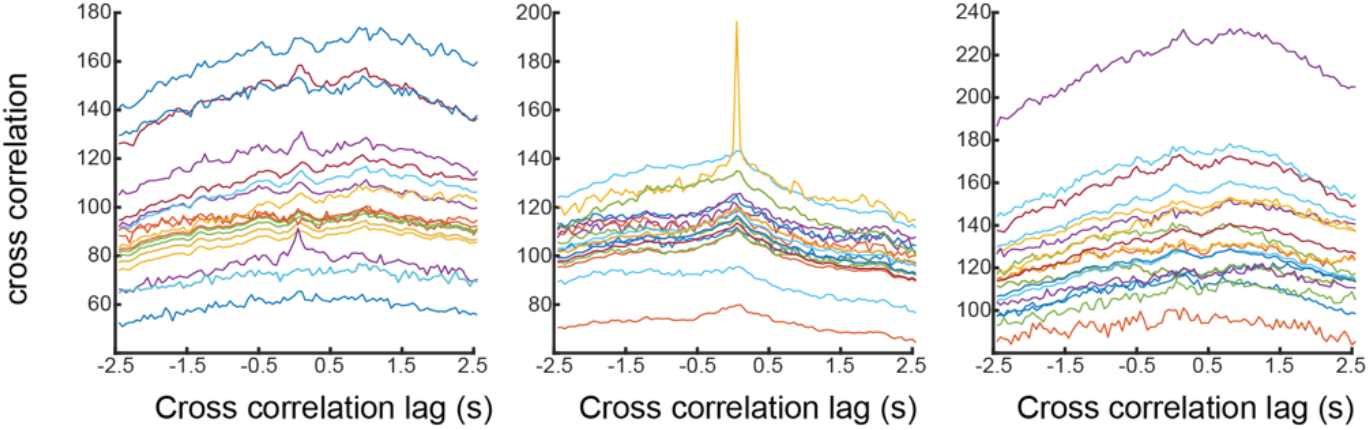
Raw cross correlation values between the full DF/F0 trace of three example events with the full DF/F0 traces of 20 random events from the same video. The lack of a clear peak in the majority of cross correlations shows that the majority of events are not correlated in time.

**Supplemental Figure 3:**
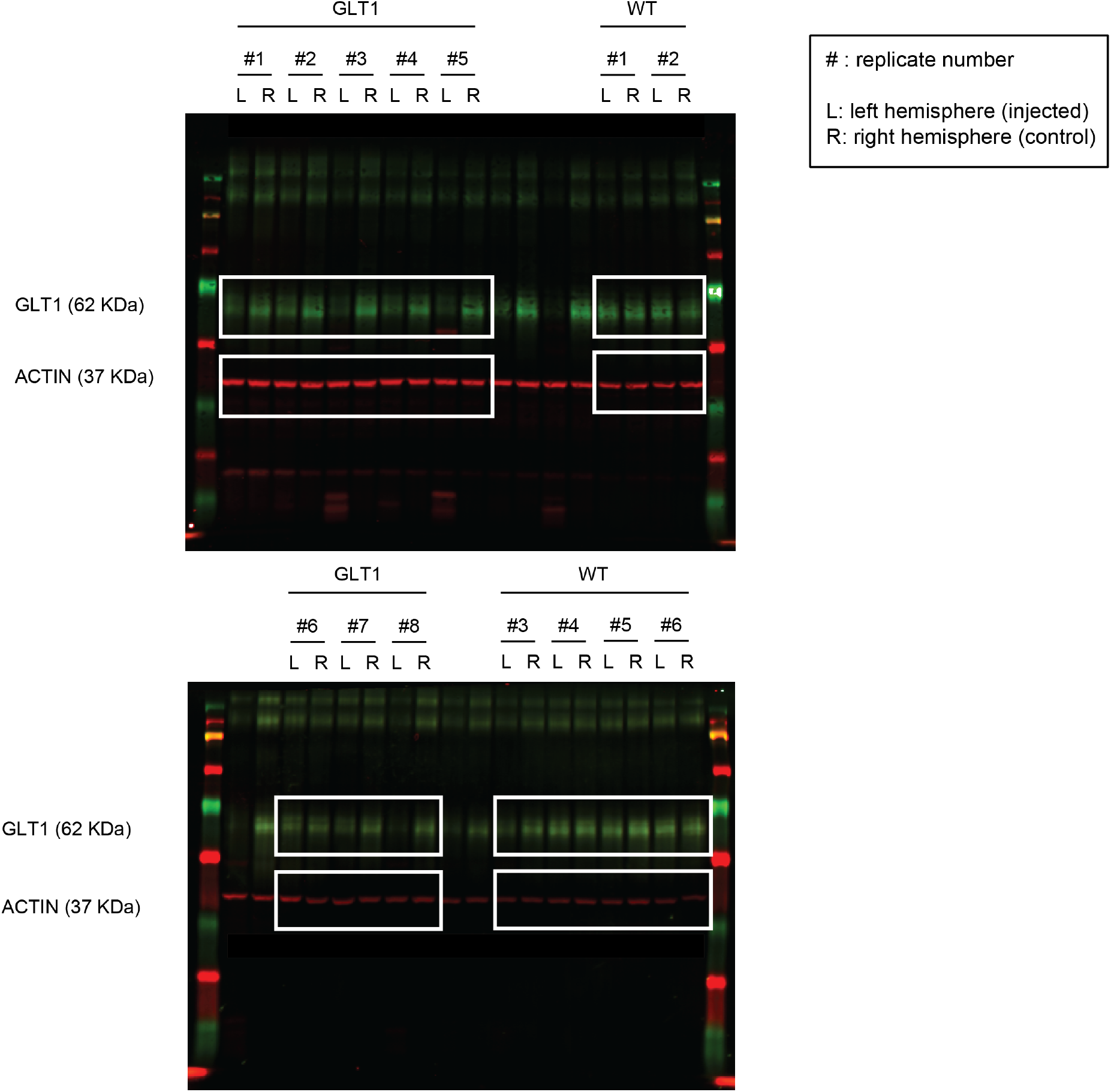
Annotated raw images of Western Blot. White boxes indicate samples used for quantification of GLT1 protein levels described in Figure 3. Each replicate is a single mouse, M1 cortex from left (L) and right (R) hemispheres were dissected and protein levels measured by Western Blot.

**Supplemental Figure 4:**
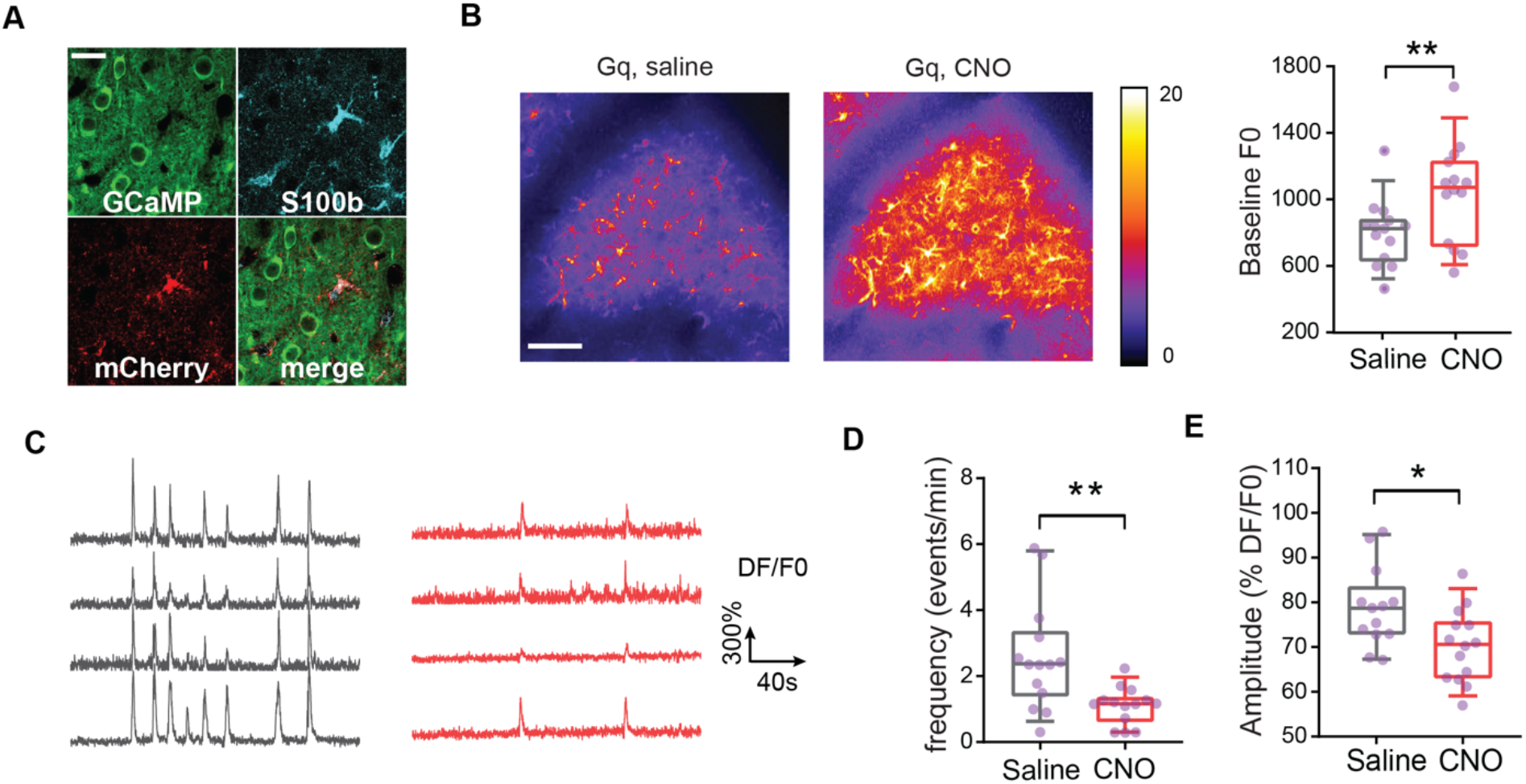
Calcium Activity in Gq Activated Astrocytes. **A.** h3MD(Gq)-mCherry colocalized with immunohistochemistry labeling of astrocyte marker S100b but not with neuronal GCAMP. Scale bar, 25μm. **B.** Gq activation increased the levels of cytoplasmic calcium. The same field-of-views containing AAV-GFAP-h3MD(Gq)-mCherry expressing astrocytes (“Gq”) in naïve (untrained) GFAP-GCaMP5G mice were imaged over 10 min passive sessions, 24 hours apart, and 30 min after IP injection of vehicle (saline) or clozapine-N-oxide (CNO). **Left:** Colormaps of the projection of the average GCAMP fluorescence (color bar) in astrocytes, in a 10 min imaging session. Example astrocytes GCaMP fluorescence imaging sessions (24 hours apart) of the same field of view, 30 min after IP injection of either clozapine-N-oxide (CNO) or vehicle (saline). Scale bar represents 100 µm. **Right:** Quantification of calcium baseline levels (average GCAMP fluorescence) from the three imaging sessions. (n= 14 non-overlapping fields of view from 5 mice, Gq+saline: mean = 802.8±53.07, Gq+CNO: mean= 1043±80.06, **: p=0.0093, paired t-test). **C.** Example astrocyte ROI raw DF/F0 traces (black, Gq+saline; red, Gq+CNO). **D.** Frequency of spontaneous calcium events (n= 14 non-overlapping fields of view from 5 mice, Gq+saline: mean = 2.596±0.4362, Gq+CNO: mean= 1.11±0.1487, **: p=0.0045, paired t-test). **E.** Quantification of the average event amplitude (DF/F0) (n= 14 non-overlapping fields of view from 5 mice, Gq+saline: mean = 78.87±2.490, Gq+CNO: mean= 70.26±2.178, *: p=0.0357, paired t-test). Box plots as defined in Fig. 2B.

**Supplemental Figure 5:**
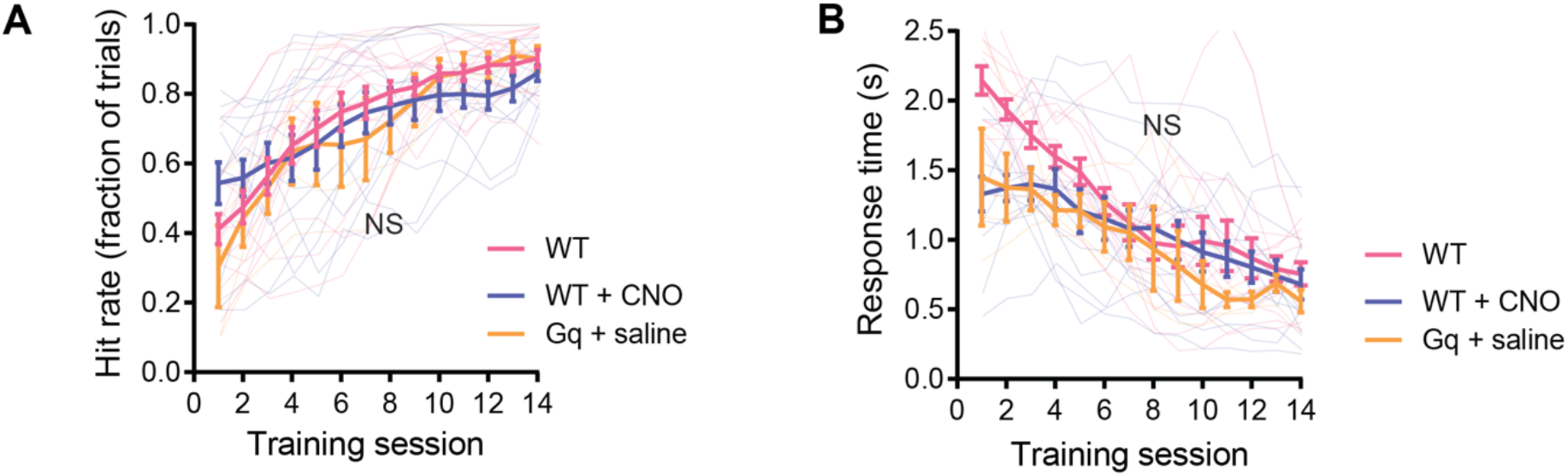
Gq-DREADD expression in M1 astrocytes does not have a CNO-independent effect on hit rate and response time. **A.** Hit rate (n= 14 WT, 12 WT + CNO, 4 Gq +saline mice, NS: p=0.8382 for variation explained by group, p<0.0001 for variation explained by training time, two-way Repeated Measures ANOVA). **B**. Response time (n= 14 WT, 12 WT + CNO, 4 Gq +saline mice, NS: p=0.0809 for variation explained by group, p<0.0001 for variation explained by training time, two-way Repeated Measures ANOVA).

**Supplemental Figure 6:**
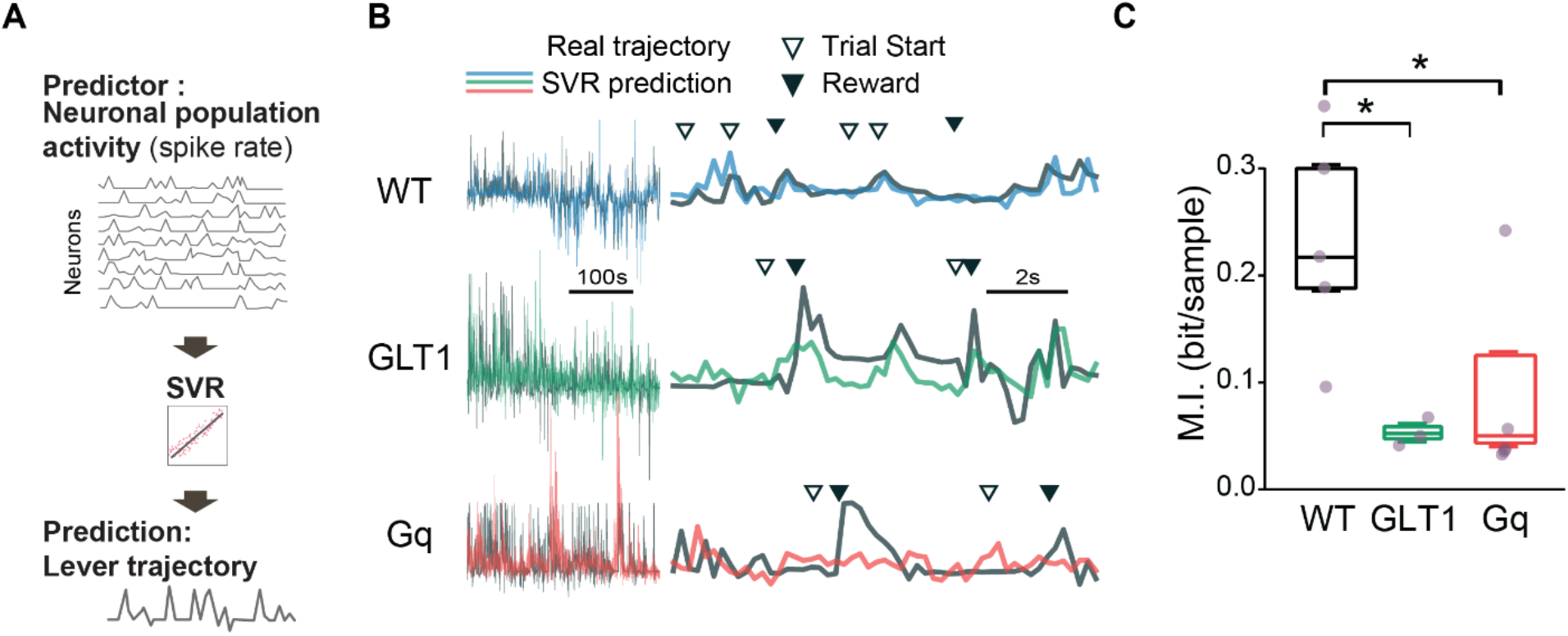
M1 neuronal population decoding of movement trajectories. **A.** A support vector regression (SVR) decoding model was used to predict the push trajectory during each training session from neuronal population spiking rate. **B.** Example lever movement traces decoded from the neuronal population activity, compared to actual traces. Black: actual movement trajectory; Blue: decoded trace from WT animals; Green: decoded trace from GLT1 animals; Red: decoded trace from Gq animals. **Left**: Predicted and actual traces; scale bar represents 100s. **Right:** zoomed in traces; scale bar represents 2s. Arrows indicate trial start (white) and reward or time when lever position reached reward threshold (black). **C**. Decoding performance (mutual information) was decreased in both GLT1 and Gq mice. N = 5/3/5 mice for WT/GLT1/Gq respectively; WT: median=0.219, GLT1: median=0.054, *: p = 0.0357, Mann-Whitney U test; WT: median=0.219, Gq: median=0.052, *: p=0.0318, Mann-Whitney U test. N= 13/11/16 non-overlapping fields of view, 33 to 91 neurons per field-of-view, total 580/586/565 neurons and 734/911/631 trials, from 5/3/5 WT/GLT1/Gq mice respectively. Box plots as defined in Fig. 2B.

**Supplemental Figure 7:**
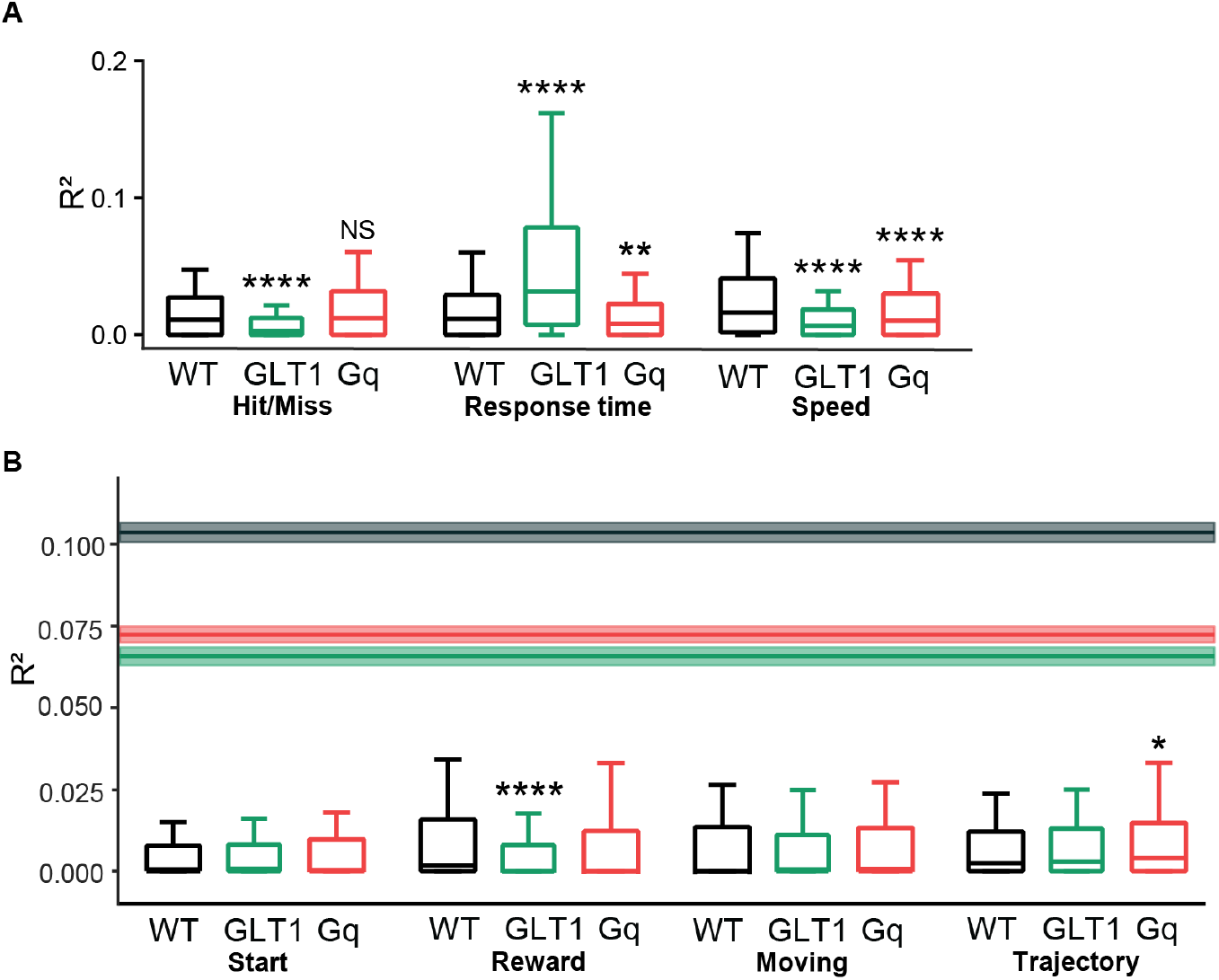
Encoding Performance of Behavioral Predictors for Neuronal Activity. **A.** Encoding performance of the GLM, as described in Fig.8, for the behavioral features with R^2^>1%: Hit/Miss, response time and speed. GLT1 mice showed less encoding of the trial outcome Hit/Miss (****: p=1.17E-17, Mann-Whitney U test) whereas Gq mice showed no significant difference from WT (NS: p=0.208, Mann-Whitney U test). Response time was encoded less in Gq mice (****: p=0.00201, Mann-Whitney U test) but more in GLT1 mice (****: p=1.38E-22, Mann-Whitney U test), consistent with the behavioral differences observed in these mice. Both astrocyte manipulations reduced push speed encoding in M1 neurons (WT vs Gq, ****: p=7.01E-6; WT vs GLT1, ****: p = 2.62E-17; Mann-Whitney U tests), consistent with the disruptive effect of Gq and GLT1 manipulations on push trajectory decoding. **B.** Encoding performance of other behavioral features, with R^2^<1%: start, reward, lever move and lever trajectory. Horizontal lines show the median predictive power of the full model in 3 experimental groups, and the shaded areas show their respective notch size. Neurons in GLT1 animals showed less encoding of the movement onset/reward (WT vs GLT1, ****: p=4.30E-7, Mann-Whitney U test). Gq neurons showed a moderately higher encoding of push trajectory than WT (WT vs Gq, *: p = 0.0482, Mann-Whitney U test). N= 13/11/16 non-overlapping fields of view, 33 to 91 neurons per field-of-view, total 580/586/565 neurons and 734/911/631 trials, from 5/3/5 WT/GLT1/Gq mice respectively. Box plots as defined in Fig. 2B.

### Supplemental Tables 1-3

**Supplemental Table 1: List of Differential Expression Genes (DEGs) in M1 Astrocytes During Motor Learning** Table of DEGs identified using EdgeR, for both novice/naïve and expert/naïve comparison with logarithm of fold change (logFC) and p=-value (PValue). The last two columns specify if the gene is a DEG (p-value<0.05) for the novice/naïve comparison (“novice”) or expert/naïve comparison (“expert”).

**Supplemental Table 2: List of DEGs Enriched GO categories** Table of significantly enriched GO biological process, molecular function and cellular component categories, from the DEG list. P-values are specified for the novice/naïve(“novice”) and expert/naïve (“expert”) comparisons.

**Supplemental Table 3: Gene Set Enrichment Analysis in M1 Astrocytes During Motor Learning.** Significantly enriched GO biological process gene sets from the whole dataset. Number of expressed genes (N Genes) per gene set, false discovery rate (FDR) and gene symbols are specified.

### Supplemental Videos 1-2

**Supplemental Video 1: Example Calcium Imaging of M1 layer 2/3 Astrocytes During Expert Training Session. Top:** Overlay of lever position trace (white in black box), trial events (white square indicates trial start; green square indicates time when the lever passes the threshold and reward is delivered), and GCaMP6f-lck fluorescence acquired by two-photon microscopy in an awake, behaving, expert mouse. **Bottom:** Corresponding overlay colored spatiotemporal event area as detected by the AQuA analysis. 80.3 x 40.2 μm field of view. Scale bar represents 10μm. Video speed 1x, 11fps.

**Supplemental Video 2: Example Calcium Imaging of M1 Layer 2/3 Neurons During Expert Training Session.** Overlay of lever position trace (white in black box), trial events (white square indicates trial start; green square indicates time when the lever passes the threshold and reward is delivered), and GCaMP6s fluorescence acquired by two-photon microscopy in an awake, behaving, expert mouse. 274 x 274 µm field of view. Video speed 1x, 5fps.

